# Transcriptional profile of pyramidal neurons in chronic schizophrenia reveals lamina-specific dysfunction of neuronal immunity

**DOI:** 10.1101/2020.01.14.906214

**Authors:** Xiaojun Wu, Rammohan Shukla, Khaled Alganem, Erica Depasquale, James Reigle, Micah Simmons, Chang-Gyu Hahn, Vahram Haroutunian, Jarek Meller, James Meador-Woodruff, Robert McCullumsmith

## Abstract

While the pathophysiology of schizophrenia has been extensively investigated using homogenized postmortem brain samples, few studies have examined changes in brain samples with techniques that may attribute perturbations to specific cell types. To fill this gap, we performed microarray assays on mRNA isolated from anterior cingulate cortex (ACC) superficial and deep pyramidal neurons from 12 schizophrenia and 12 control subjects using laser capture microdissection. Among all the annotated genes, we identified 134 significantly increased and 130 decreased genes in superficial pyramidal neurons, while 93 significantly increased and 101 decreased genes were found in deep pyramidal neurons, in schizophrenia compared to control subjects. In these differentially expressed genes, we detected lamina-specific changes of 55 and 31 genes in superficial and deep neurons in schizophrenia, respectively. Gene set enrichment analysis (GSEA) was applied to the entire pre-ranked differential expression gene lists to gain a complete pathway analysis throughout all annotated genes. Our analysis revealed over-represented groups of gene sets in schizophrenia, particularly in immunity and synapse related pathways in pyramidal neurons, suggesting the disruption of these pathways plays an important role in schizophrenia. We also detected pathways previously demonstrated in schizophrenia pathophysiology, including cytokine and chemotaxis, post-synaptic signaling, and glutamatergic synapses. In addition, we observed several novel pathways, including ubiquitin-independent protein catabolic process. By comparing our differential expression gene profiles with 51 antipsychotic treatment datasets, we demonstrated that our results were not influenced by antipsychotic treatment of our subjects. Taken together, we found pyramidal neuron-specific changes in neuronal immunity, synaptic dysfunction, and olfactory dysregulation in schizophrenia, providing new insights for the cell-subtype specific pathophysiology of chronic schizophrenia.

## Introduction

Schizophrenia is a severe mental illness characterized by hallucinations, delusions, abnormal behavior, and disorganized speech. It is a relatively common disorder that is the seventh most costly disease to our society, and available medication treatment is only partially efficacious [1, 2]. While work to understand the causes and pathology of schizophrenia is ongoing, more effective therapeutic approaches need to be developed. We postulate that a more sophisticated understanding of the pathophysiology of schizophrenia will lead to such breakthroughs.

Our previous reviews have described the “blender” problem in postmortem brain studies, pointing out that results using blended tissue homogenates could be difficult to interpret and may lose important information when samples are a mixture of neurons, astroglia, microglia, and/or endothelial cells [3, 4]. For example, in our prior cell-subtype specific studies, we showed that the EAAT2 splice variant EAAT2b mRNA was increased in schizophrenia in anterior cingulate cortex (ACC) pyramidal neurons, but not in region-level homogenized samples, while another splice variant, EAAT2 exon9skipping, was increased at the region level, but not changed in pyramidal neurons [5-7]. These findings emphasize the limitations associated with measuring gene expression in blended human brain samples.

In this study, we targeted cortical pyramidal neurons, as they may easily be identified for capture using a modified RNase-Free Nissl stain [4-10]. Cortical pyramidal neurons form neuronal circuits in the cerebral cortex and are required to process sensory, planning, and executing functions [11]. Laser capture microdissection (LCM) is a technique frequently used to isolate specific cell types from postmortem tissues under microscopic visualization [4-10], which is a powerful tool for cell level investigations. In the current study, we isolated pyramidal neurons from the ACC of 12 schizophrenia and 12 control subjects by LCM. Neurons from superficial (lamina II-III) and deep (lamina V-VI) cortical layers were captured from each subject. Their transcriptional profiles were assessed by a microarray analysis using the GeneChip® Human Gene 1.0 ST Arrays, comprised of more than 750,000 unique 25-mer oligonucleotides representing transcripts for more than 20,000 human genes, with each gene represented on the array by approximately 27 probes spread across the full length of its transcript. Differentially altered pathways between schizophrenia and control subjects were analyzed using gene ontology databases and Gene Set Enrichment Analysis (GSEA) [12].

## Materials and methods

### Subjects

Postmortem tissue from anterior cingulate cortex (ACC) in subjects diagnosed with schizophrenia (SCZ; n=12) and a control group (NC; n=12) were obtained from the Mount Sinai/Bronx Veterans Administration Medical Department of Psychiatry Brain Bank. Schizophrenia and control subjects were matched for age of death, postmortem interval (PMI), sex, and tissue pH (Table 1). Tissue was collected in compliance with the Mount Sinai School of Medicine Institutional Review Board protocol for postmortem tissue [9].

**Table 1:** Subject demographics.

|  | Control | Schizophrenia |
| --- | --- | --- |
| N | 12 | 12 |
| Sex | 6 Male/6 Female | 6 Male/6 Female |
| Tissue pH | 6.4 ± 0.3 (6-6.82) | 6.3 ± 0.4 (5.8-7.1) |
| PMI* (Hours) | 10.2 ± 6.3 (3.38-20.4) | 11.9 ± 5.4 (6.2-20.4) |
| Age (Years) | 73.5 ± 10.8 (58-89) | 72.6 ± 10.8 (57-90) |
| *Medication | 0/12/0 | 8/2/2 |

\*Medication (On/Off/Unknown)

### Harvest of neurons from superficial and deep laminas of ACC

Fresh frozen anterior cingulate cortex from subjects was sectioned at 14 µm on SuperFrost^®^/Plus glass slides (Fisher Scientific, Pittsburgh, PA) and stored at −80°C until use. For laser-capture microdissection (LCM) [9], tissue slides were incubated in RNase-free 1% cresyl violet acetate for 2 minutes, submerged in 95% ethanol, and then in 100% ethanol for 30 seconds, followed by immersion in xylene for 5 minutes. Lamina and pyramidal neurons were defined based on distribution across the cortical thickness and morphology of cells [9, 13, 14] and approximately 2000 pyramidal neurons were isolated from superficial (lamina II-III) and deep (lamina V-VI) layers from each subject by LCM using the Arcturus VERITAS instrument and protocols confirmed by nissl staining of adjacent sections. LCM was performed under 20X objective lens and with laser settings ranging from 70-100 mW in power, and 2,000-3,000 µsec in duration. Neurons were collected onto a CapSure® Macro LCM Cap and 50µl extraction buffer from the PicoPure® RNA Isolation Kit (Applied Biosystems™) was added to the cells, followed by an incubation at 42°C for 30 min. The cell lysates were spun at 800 *× g* for 2 min and the supernatant was stored at −80°C until use. We have previously reported in multiple studies that we are capable of isolating samples enriched for pyramidal neurons [4-10].

### RNA isolation, cDNA synthesis, labeling, and oligonucleotide array hybridization

Total unpooled RNA from the 2000 pyramidal neurons from each subject was purified with the QuickGene RNA Tissue kit SII in conjunction with the QuickGene Mini-80 semi-automated station (FujiFilm Life Sciences Division). RNA samples were processed by Genome Explorations Inc., Memphis, TN – a certified Affymetrix® microarray service provider – for microarray assays. Immediately prior to cDNA synthesis, the purity and concentration of RNA samples were determined by readings of OD_260/280_ using a dual beam UV spectrophotometer and RNA integrity was determined by capillary electrophoresis using the RNA 6000 Nano Lab-on-a-Chip kit and the Bioanalyzer 2100 (Agilent Technologies, Santa Clara, CA) following the manufacturers instructions. The purified RNA was amplified with the WT-Ovation™ Pico RNA Amplification System (NuGEN Technologies, Inc). 5-20 ng RNA from each sample was reverse transcribed to complementary DNA (cDNA), amplified, fragmented, and labeled with biotin using the Ovation Pico WTA, Ovation Exon Module, and Encore Biotin Module kits according to the manufacturer’s instructions (NuGEN, San Carlos CA).

### Microarray assay

The GeneChip® Human Gene 1.0 ST Arrays (Affymetrix®, Santa Clara CA) were used. Arrays were washed and stained with phycoerythrein-conjugated streptavidin (Life Technologies, Carlsbad, CA) in a Fluidics Station 450 (Affymetrix) according to the manufacturer’s recommended procedures. Fluorescence intensities were determined using a GCS 3000 7G high-resolution confocal laser scanner.

### Differential expression analysis and pathway enrichment analysis

To identify differential expression genes and biological pathways, Limma (Linear Model of Micro Array) R package and gene set enrichment analysis (GSEA) were utilized. Full details are included in the supplement.

### Antipsychotic analysis

Antipsychotics may have effects on gene expression [15, 16]. Therefore, we compared our data with 51 antipsychotic treated profiles. We used tmod package in R to measure the area under the receiver operating characteristics (AUROC) curve. The method uses ranked genes associated with different contrasts to measure the enrichment of up and down regulated gene signatures from different antipsychotic drugs tested in the mouse and rat [17-21] and computes p-value based on CERNO statistics [22]. Datasets GSE4031, GSE66275, and GSE89873 were also included in our antipsychotic analysis and have not been published in previously reported studies to the best of our knowledge. The AUC (Area Under the Curve) values calculated is equal to the probability that an antipsychotic associated gene signature will rank higher in schizophrenia associated ranking than the signature not enriched for a given antipsychotics. The AUC scores calculated for ranked genes (separating upregulated and downregulated genes) associated with an antipsychotic against the signature of another similar antipsychotic is expected to show a higher value and can be used as a positive control for our analysis.

### In silico confirmation study

The top 100 differential expression genes with the highest significance (lowest *P value*) from superficial, deep, and interaction of superficial and deep were selected. The overlapping genes among the three analyses were picked and the noncoding RNA genes removed. 56 genes that met our criteria were used for following analysis. We applied the selected genes in Kaleidoscope (https://kalganem.shinyapps.io/BrainDatabases/) to determine the similarity of our analysis and previous reported datasets in schizophrenia.

## Results

### Altered gene expression profile in schizophrenia

Postmortem tissues were obtained from 12 schizophrenia patients and 12 psychiatrically healthy comparison subjects. 2000 pyramidal neurons were enriched from both superficial and deep layers of the anterior cingulate cortex of each subject by LCM (96,000 cells total) (Fig. 1A, Table 1). The RNA of each LCM sample was purified, reverse transcribed, and 48 separate arrays were performed, one for each superficial or deep samples for each subject.

**Figure 1.**
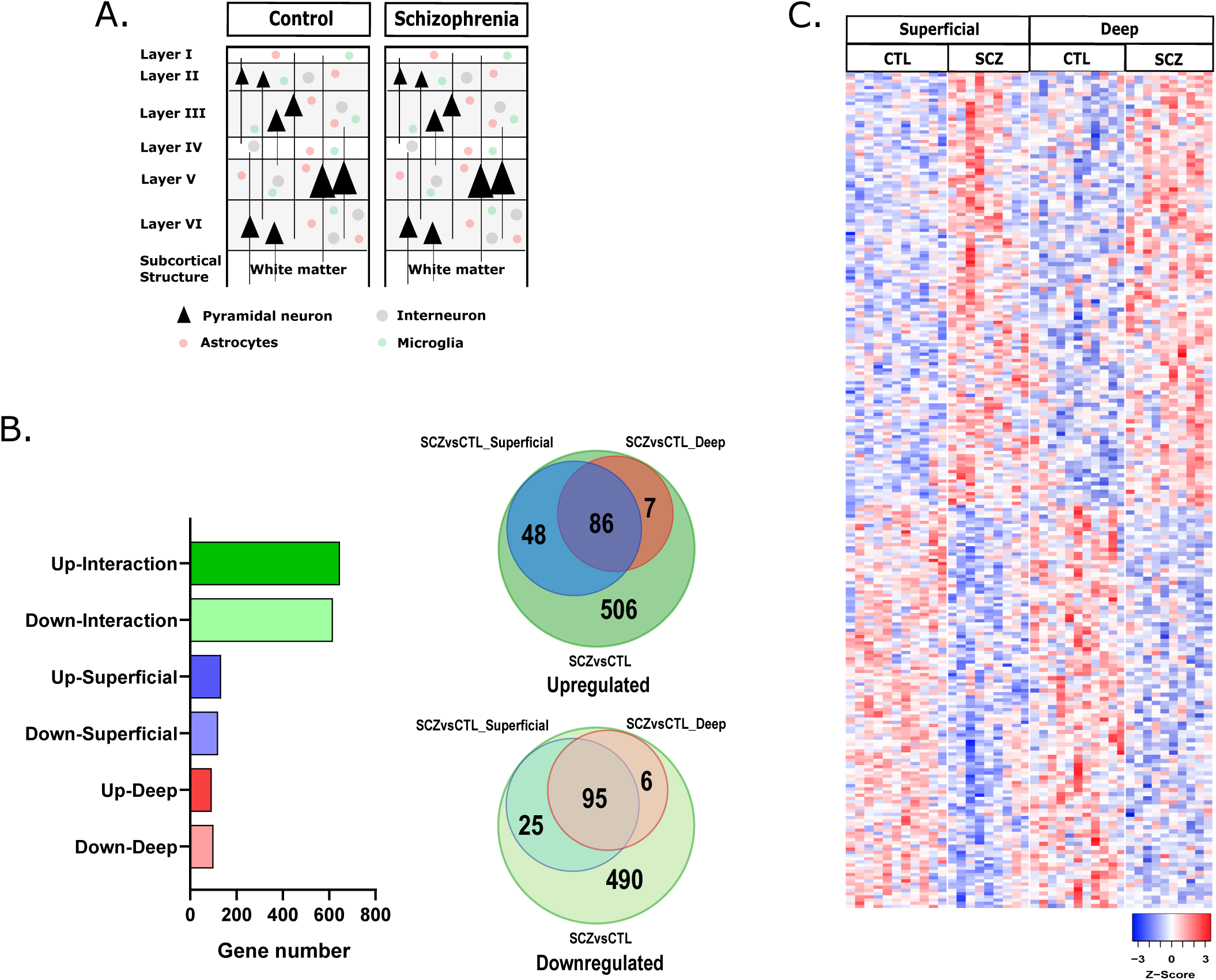
Distinct gene expression profiles detected in schizophrenia patients. A. Superficial (Layer II and III) and deep (Layer V and VI) pyramidal neurons were enriched from anterior cingulate cortex of schizophrenia and control subjects. B. Venn diagrams show differential expression gene numbers overlapping in superficial and deep neurons to the comparison groups regardless of layers between schizophrenia and control subjects. C. Differentially expressed genes in the superficial and deep neurons were shown by heatmap, using www.heatmapper.ca. Clustering method: complete linkage; Distance measurement method: pearson. Plotted genes of *P* < *0.05*.

Differential expressed genes were first analyzed to compare the schizophrenia and control subjects. A total of 22,595 genes were annotated from the arrays. Among which, we identified 134 significantly increased and 130 decreased genes in schizophrenia superficial neurons, while 93 significantly increased and 101 decreased genes were found in schizophrenia deep neurons, compared to control subjects (*P* < *0.05*, Fig. 1B). Among these genes, 55 genes were exclusively differentially expressed in the superficial neurons, while 31 genes were exclusively differentially expressed in the deep neurons. In the comparison of combined schizophrenia superficial and deep neurons, we identified 647 significantly increased and 616 decreased genes, compared to control subjects (*P* < *0.05*, Fig. 1B). Interestingly, these 1263 differentially expressed genes include all the increased and decreased genes in the superficial and deep neurons, although these differentially expressed genes from the superficial and deep neurons do not completely overlap (Fig. 1B). Based on unsupervised clustering, the schizophrenia and control subjects clustered well and the significantly increased and decreased differentially expressed genes are clearly separated between the groups (Fig. 1C). The top 50 up and down regulated genes are listed in Supplementary Table 1.

### Pathway alterations involved in schizophrenia pathology

We performed gene set enrichment analysis (GSEA) with the complete list of the annotated genes ranked based on their expression and significance levels. GSEA revealed multiple gene sets involved in schizophrenia, including proteolysis, immunity, synapses, protein modification, receptors, transporters, and sensory system (supplementary Table 2). Superficial and deep neurons share 45% of the pathways, including upregulated and downregulated gene sets. However, they have opposite directionality for the remaining 55% pathways, including proteolysis, antigen processing and presentation, and axons (Supplementary Table 3). This result suggests that schizophrenia may affect the superficial and deep neurons differently.

#### Synapse

In our microarray data, 13 synapse-related pathways were identified, with 9/13 involved in postsynaptic signals and 8/9 involved in postsynaptic density and plasma membrane. There were no downregulated synaptic pathways detected. 6/9 synaptic pathways upregulated in deep neurons were also upregulated in superficial neurons. 5/6 of these pathways are related to postsynaptic functions (Supplementary Table 2).

#### Proteolysis

While most gene set categories were detected in both superficial and deep neurons, four were detected only in the deep neurons, including the proteolysis pathway. The proteolysis pathways detected in our data analysis is mostly affected in deep (7/8 pathways), but not superficial, neurons. They are mostly involved in ubiquitin-dependent and -independent protein catabolic processes, modification-dependent and proteasomal protein catabolic processes (Supplementary Table 2).

#### Protein modification

Protein modification pathways linked to mitosis, protein folding, protein ubiquitination, oxidation, and phosphorylation (Supplementary Table 2) were also altered in our study. Protein folding and oxidation pathway alterations were as observed in the superficial neurons, while ubiquitination and ERK pathways are hits in the superficial and deep neurons.

#### Immunity

Our microarray data shows strong correlation between schizophrenia and immunity. The immunity gene set hits in our analysis are mostly downregulated in the schizophrenia group. 96% and 89% of detected immunity gene sets were downregulated in the superficial and deep neurons, respectively. When comparing schizophrenia and control subjects combining superficial and deep neurons, 89% of the enriched gene sets were downregulated. When we looked at the gene set categories, we found that 54% gene sets are cytokine and chemotaxis related, including Il-8, IL-10, and TNFα regulatory pathways. (Fig. 2).

**Figure 2.**
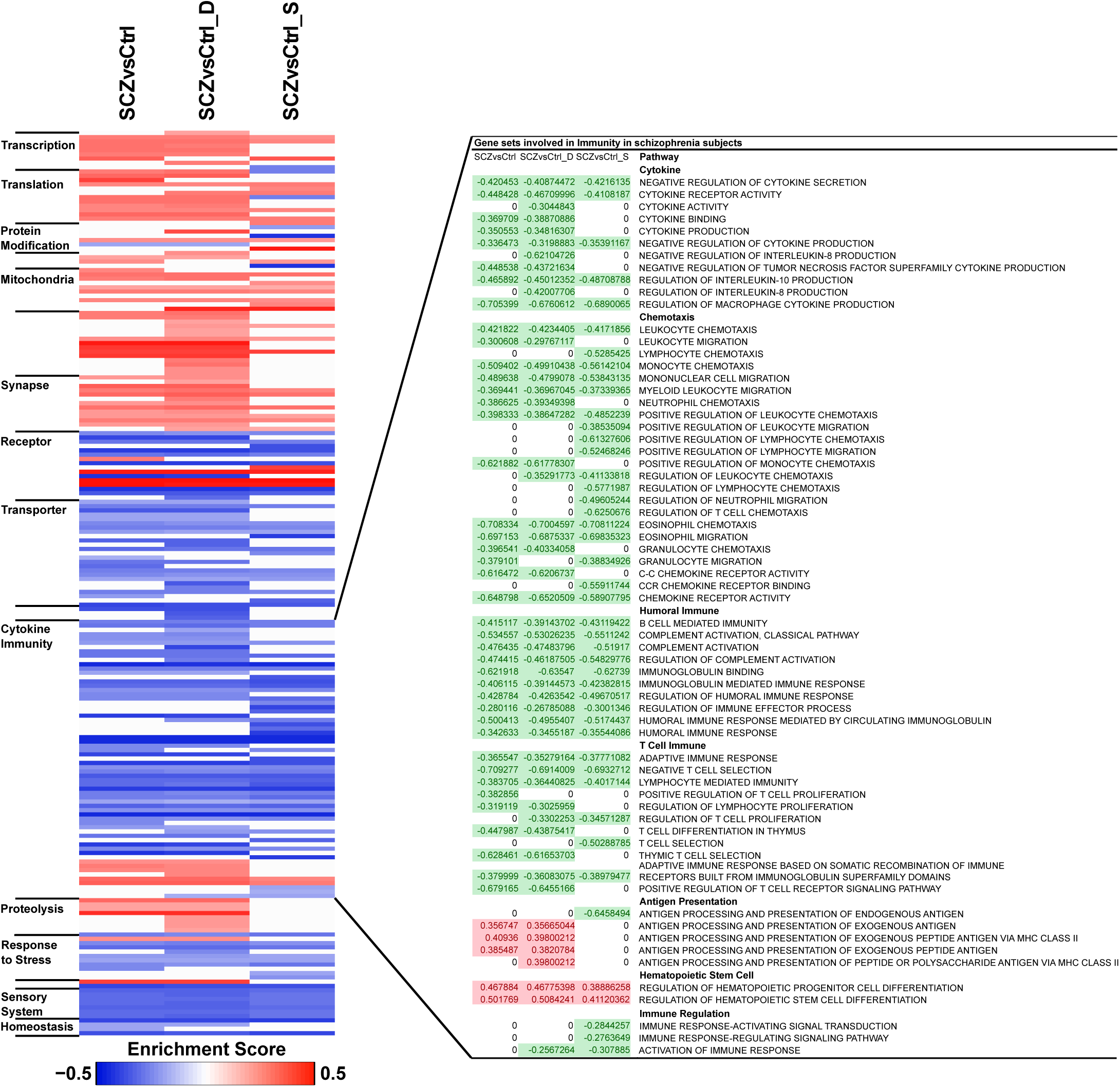
Gene set enrichment analysis reveals pathways involve in schizophrenia pathophysiology. Differentially expressed genes from each comparison were ranked based on their expression value and significance. GSEA was performed using the pre-ranked gene lists. Heatmap was generated based on the enrichment score and significance of GSEA (*P* < *0.05*). Cytokine immunity pathways were enlarged to show details.

Humoral immunity gene sets, including B cell mediated immunity, complement, and immunoglobulin mediated immunity, were hits in our gene set enrichment analysis, T cell activation pathways are also involved, indicating the involvement of adaptive immunity. Interestingly, there are different patterns between superficial and deep in certain gene sets. For example, we found 5 antigen presentation related gene sets total and 4 of them are upregulated in the deep neurons only while 1 is downregulated in the superficial neurons only. The cytokine IL-8 and TNFα gene sets are dysregulated only in the deep neurons, but not in the superficial neurons.

### Antipsychotic medication

In order to determine whether the pathway alterations we have observed were driven by antipsychotic medication, we compared our differential expression profiles to 51 antipsychotic drug treated transcriptional profiles from rodents (Supplementary Table 4). We found that the profiles of deep neuron and the total (superficial and deep neuron combined) have no significant similarity with any of the antipsychotic drug profiles in either up or down direction (shown in grey curves, Figure 3). However, the superficial neurons showed subtle (AUC of 0.55) but significant (*adj. P. value* < *0.05*) similarity with 3/51 downregulated profiles associated with: GSE66275: Risperidone, 5mg/kg/d, 21d; GSE66275: Haloperidol, 0.25mg/kg/d, 21d, and GSE48955: Haloperidol, 1mg/kg, 8h (Figure 3 green curves), and 1/51 upregulated profile associated with GSE677: Haloperidol, 3mg/kg/d, 30d (Figure 3 red curve). Note that the AUC values of these datasets are less than 0.55, while 0.5 means no similarity.

**Figure 3.**
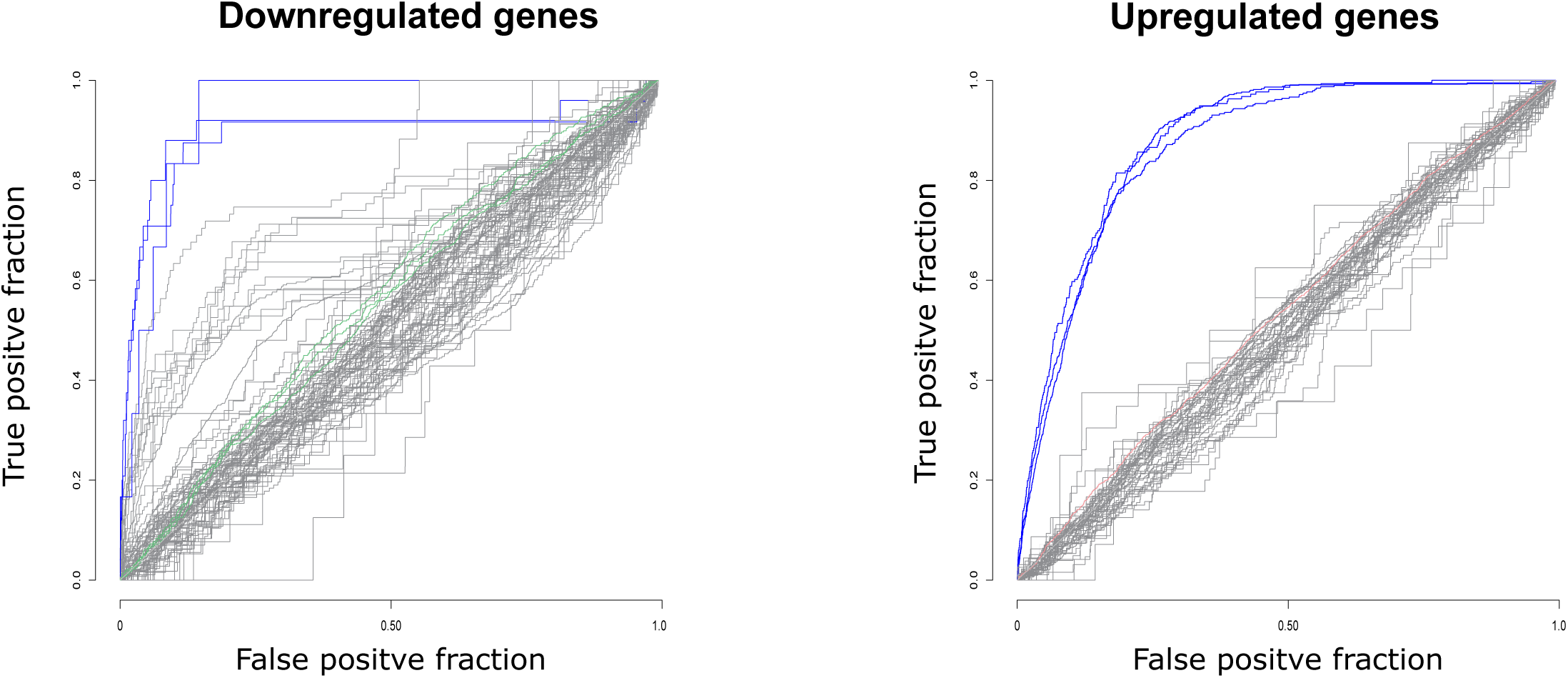
Our results are not influenced by antipsychotic drugs. 51 previous published antipsychotic drug profiles were compared with the 3 indicated comparisons of the current study. Each line represents 1 antipsychotic profile. The profiles with significant similarity with our data are either marked in red (based on ranked upregulated genes) or green (based on ranked downregulated genes). Grey curves represent the non-significant profiles. Blue curves represent the significant positive control profiles compared to the clozapine consensus with AUROC higher than 0.85. (*adj. P* < *0.05*)

### Confirmation studies

To compare our study with previous datasets, we selected the overlapping top differentially expressed genes from our analyses of superficial, deep, and superficial plus deep, yielding a total of 56 genes. We applied the selected genes in the R shiny App Kaleidoscope (https://kalganem.shinyapps.io/BrainDatabases/) to evaluate the similarity of their expression between our analysis and previously reported datasets (Fig. 4). Our results are well clustered with the schizophrenia datasets in Kaleidoscope. In the cell level comparisons, the superficial and deep neuron profiles are clustered best with the human iPSCs of neurons, NPCs, and DLPFC layer 3 and layer 5 pyramidal neurons (Lewis_2015_L3, Lewis_2015_L5, Lewis_2017_L3, and Lewis_2017_L5).

**Figure 4.**
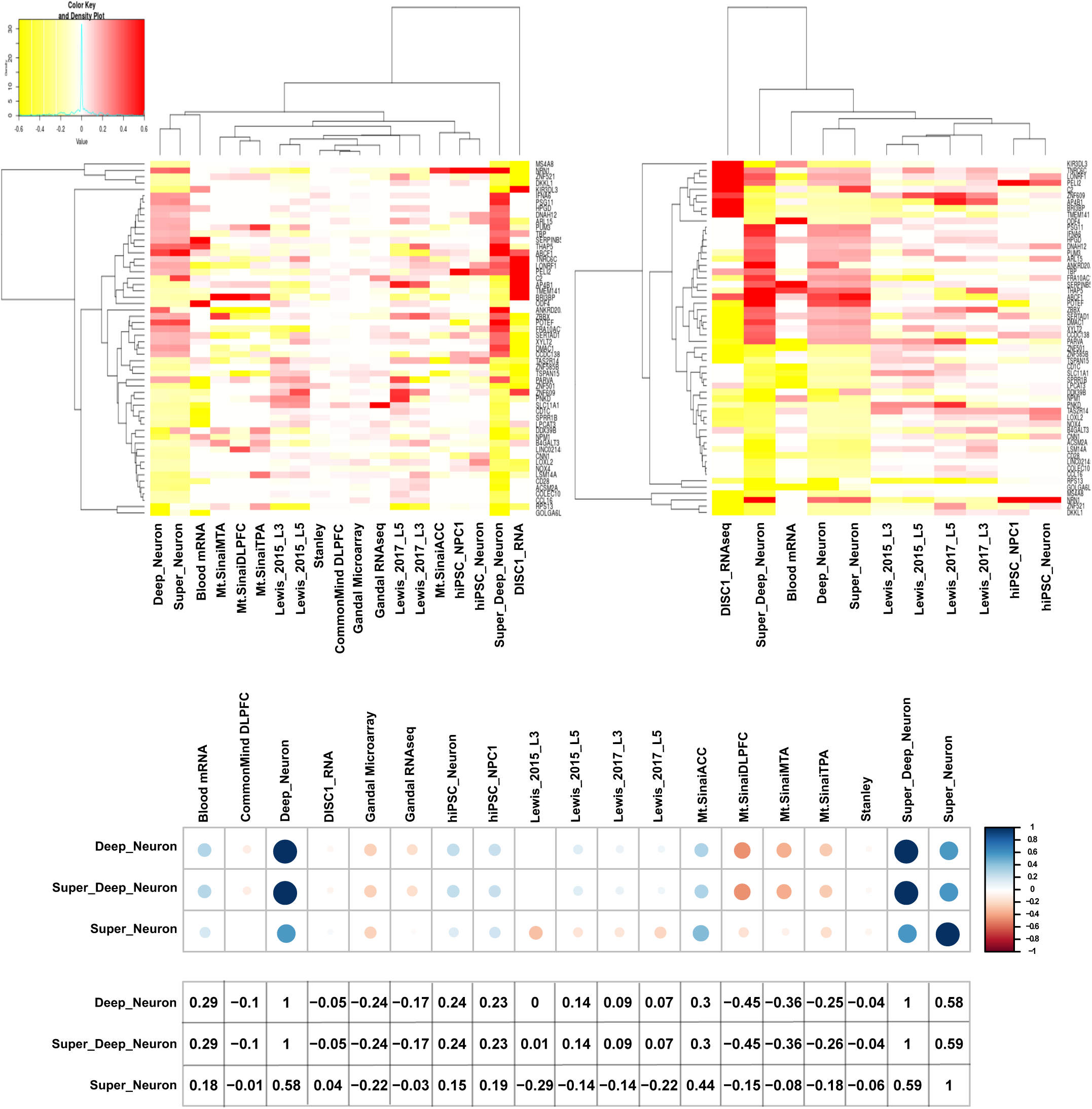
Current microarray analysis shows positive concordance score with multiple previously reported schizophrenia datasets. The top 56 overlapping genes from all three comparisons were selected and used to compare with previously reported schizophrenia datasets in the R shiny App Kaleidoscope. Concordance scores were calculated based on the Log_2_ fold change. Red represents negative correlation and blue represents positive correlation. The color intensity and circle sizes represent the levels of concordance (Please see supplementary materials for detailed description).

## Discussion

A previous microarray analysis in the pyramidal neurons in the DLPFC laminas III and V separately found that schizophrenia patients had robust pathway alterations in mitochondrial function and ubiquitin proteasome system [23]. A more recent microarray study in the same cortical layers and cell types using a different pathway analysis methods (INGENUITY® Pathway Analysis) showed that the most striking pathways involved in oxidative phosphorylation and mitochondrial dysfunction [24]. While these pathways were also detected in our microarray analysis (Supplementary Table 2), they are not in the most remarkably altered pathways. These differences may be due to 1) the subjects in our study have different age, disease length, and medication history, 2) different brain regions were utilized (ACC *vs*. DLPFC) [25], 3) normalization, differential expression analysis approaches and statistical analysis differences may impact the differential expression genes, and/or 4) our use of more cutting-edge pathway analysis methodology. They used either IPA or GO, KEGG, Biocarta and Reactome databases with only the significant differentially expressed genes. In contrast, we performed GSEA with the entire ranked gene profile, which has the advantage to detect more subtle changes in the data sets [12]. GSEA is particularly useful for human studies, which typically have more variation among subjects and thus reduced power to detect effects with analyses that only use the top ranked changes.

It’s been well documented that schizophrenia pathophysiology is partially related to immunity system, namely pro-inflammatory cytokine secretion, T cell activation, autoantibody production, and microglia dysfunction [26]. In our GSEA analysis, the most dramatically altered pathways are immunity related. Among these, 57% of them are cytokine and chemokine related, indicating cytokine and chemokine may play an important role in schizophrenia pathophysiology. Previous studies have shown alterations of cytokines TNFa, IL-6, IL-8 in schizophrenia postmortem brain in the prefrontal cortex [27], which are consistent with our results (Fig. 2). In addition, in peripheral blood mononuclear cells (PBMCs) and plasma, alterations of CCL5, MCP-1, IL-8, IL-18, IFN-r, IL-2, MIP1a, IL-1b, and IL-6 were detected [28, 29]. Although detection of these pathways is possibly partially due to cross-talk of pathways, observing these pathways clearly show the involvement of cytokines and chemokines in schizophrenia pathophysiology in pyramidal neurons. The dysregulation of cytokine IL-8 and TNFα gene sets only in the deep neurons, but not in the superficial neurons, indicates the deep and superficial neurons may possess differential regulation in certain immune pathways.

In addition to cytokines and chemokines, adaptive immune cells such as T cells and B cells activation pathways are also hits in our analysis. By combining multiple single nucleotide polymorphism (SNP)-based genome-wide association studies in schizophrenia, several significant association signals were found in the major histocompatibility complex (MHC) region on chromosome 6p21.3-22.1 [30]. This indicates involvement of adaptive immunity, especially T cells, as T cells rely on MHC molecules for selection and activation during normal and/or pathological development and priming. It has been reported that circulating T regulatory cells have increased numbers in schizophrenia subjects [31]. T regulatory cells play an important role in immune suppression, which may partially explain the overall downregulation of immune profiles in schizophrenia subjects in our GSEA analysis. Interestingly, while a majority of immune pathways are downregulated in schizophrenia, we detected upregulated antigen presentation pathways, which could be a compensation mechanism. Although the genetic association between MHC and schizophrenia is robust, it is still lacking direct experimental evidence to discover the true connection between MHC changes and schizophrenia.

The functions of B cells and autoantibodies in schizophrenia patients is somewhat controversial [32]. There is established evidence for N-methyl-d-aspartate receptor (NMDAR) hypofunction as one of the key contributors for schizophrenia symptoms [33, 34]. Autoantibodies against NMDAR were detected in a group of schizophrenia patients at early onset [35]. It is also known that diagnosis with schizophrenia has a 50% lifetime prevalence for developing an autoimmune disease [36, 37]. Autoimmune disease patients also have 55% higher chance to develop psychotic disorders [38]. However, there is no enrichment in immune loci outside of the MHC region in schizophrenia patients, which is different from several well-established autoimmune disorders [39]. Therefore, determining whether schizophrenia could be defined as an autoimmune disease requires further investigation. Our microarray data further pointed out the importance of investigating the correlation between immune dysfunction in schizophrenia and provided novel insightful knowledge specifically in pyramidal neurons.

Other than immunity related pathways, we expected to find pathways related to synapse and proteolysis, which have previously been reported in schizophrenia. Gray matter reduction in the frontal cortex of schizophrenia patients has been demonstrated by histopathological imaging [40] and this gray matter reduction is associated with synaptic loss, instead of neuron loss [41]. In addition, glutamate levels in the pre- and post-synapse between neurons and astrocytes are tightly regulated to maintain the normal synaptic functions, while glutamate dysregulation plays a critical role in psychotic diseases, such as schizophrenia [14, 42, 43]. These studies suggest the synaptic function is disrupted in schizophrenia. We detected only upregulated synaptic pathways, which may be a compensatory mechanisms. Two of the synaptic pathways upregulated only in deep neurons are asymmetric synapse and glutamatergic synapse pathways, which indicates that excitatory neurons may be more affected.

Protein degradation systems are critical for removal of damaged or toxic proteins and play an important role in normal cell homeostasis. Protein degradation dysregulation, especially ubiquitin mediated proteolysis, is associated with schizophrenia in both postmortem brains and peripheral samples [23, 44-46]. In our GSEA results, we detected both ubiquitin-dependent and – independent pathways. More interesting, 7/8 gene set hits in our analysis are upregulated in the deep neurons only. A microarray study performed analysis in pyramidal neurons from layer III and V in DLPFC also detected dysregulated ubiquitin-proteasome system in their deep neurons [23], which is consistent with our findings. There is no sufficient explanation in the literature to determine why the ubiquitin-dependent pathways are more affected in the deep schizophrenia neurons. Additional cell level investigation is required to pursue this hypothesis.

Antipsychotic medications heavily impact gene expression studies and most of our schizophrenia subjects were on typical antipsychotics (7/12). For postmortem studies, they can interfere with the interpretation of the data and with studies with small sample size it is challenging to take them in to account. To ensure that the dependent measures in our analysis are not influenced by the medication history, we compared the fold-change rank ordered transcriptional profiles of schizophrenia versus control subjects to 51 previously published antipsychotic drug treatment profiles. We only detected marginal (AUC < 0.55) similarity in superficial neurons for Risperidone, 5mg/kg/d, 21d; Haloperidol, 0.25mg/kg/d, 21d; Haloperidol, 1mg/kg, 8h and Haloperidol, 3mg/kg/d, 30d. Considering that our positive control analysis showed high similarity (AUC > 0.85, Supplementary Information) between clozapine treated profiles and the consensus treatment profile, the similarity between our results and the antipsychotic treatment profiles is minor. In addition, compared to the generally lifelong treatment of schizophrenia subjects, the antipsychotic treatment in these 4 profiles are much shorter (1 is 8h; 2 are 21d; and 1 is 30d). Our data had no significant similarity to the longer treatments (up to 12 weeks) in the profiles. This novel analysis, the first of its kind combining data for 51 drug treatment studies, using full RNA profiles, suggests our GSEA findings are not secondary to effects of treatment with antipsychotic medications.

As a confirmatory approach, we compared our top differentially expressed genes with previously reported datasets at both cellular and regional levels through an unsupervised heatmap, which showed better clustering of our datasets with cellular level studies, especially neuron and progenitor neuron iPSCs. This is expected, because different cell types possess distinct transcriptional profiles and region level datasets are consisted with a mixture of multiple cell types. However, even among the cell level datasets, we still detected genes with opposite directionality, which may be caused by subject variation, cell subtype differences, and distinct data analysis approaches. In addition, iPSCs may only partially represent native neuronal transcriptomes, as tissue culture may not reflect the *in vivo* environment. DISC1 cells represent a particular type of schizophrenia and may not have exactly the same transcriptional profile with sporadic cases of schizophrenia. We surprisingly observed high concordance score between our datasets and mRNA from blood samples [47] (Figure 4, Supplementary Results), indicating the potential of using blood samples to predict the gene expression in pyramidal neurons in schizophrenia.

There are several potential limitation for this study. While cell level studies empower the specificity of investigations to reveal transcriptional profiles of neurons in particular cortical layers, one limitation of LCM is the possible contamination of astroglia due to their close association with neurons. We have previously demonstrated robust enrichment of pyramidal neurons with our protocol, using cell-characterized neurochemical markers [5-10]. Even though microarray technology is powerful enough for many investigation purposes, it still has its limitation compared to RNA sequencing. For example, it can only detect the genes with given probes in the array, while RNA sequencing has the capacity for unsupervised gene mapping, which potentially can detect every transcript in a sample. The subject number is another limitation of this study, a necessary limitation as cutting thousands of cells is time-consuming and costly.

In summary, we compared the transcriptional profiles of superficial and deep neurons from control and schizophrenia postmortem brain lamina II and III and lamina V and VI, respectively. Our pathways analysis showed results consistent with previous reports, such as pathways involved in proteolysis, and synapse. We also had novel findings in protein modification and olfactory sensation related pathways. The most striking pathways we detected are immunity related, which further implicates the immune system in schizophrenia pathophysiology. Our study provides new information regarding the pathophysiology of schizophrenia at the cell level, yielding new directions for future investigation.

