## Supplementary information for "Transcriptional profile of pyramidal neurons in chronic schizophrenia reveals lamina-specific dysfunction of neuronal immunity"

### Supplementary Introduction

#### *Function of cortical neurons and ACC in cognition*

The superficial neurons reside in cortical layers II and III. They generally receive cortico-cortical associative projections and form synaptic connections with cerebral nuclei, such as claustrum, amygdala, and basal ganglia, and integrate sensory and motor elements [1]. The deep neurons are in cortical layers V and VI. They project to other layers within their cortical columns and also project into subcortical structures [1, 2]. Since superficial and deep neurons have not previously been investigated in the same study, we evaluated them separately [3].

The ACC plays an important role in various cognitive functions, such as response monitoring, error detection and processing, and implementing optimal strategies [4-6]. Changes in the ACC are associated with multiple cognitive disorders, including schizophrenia [7-9]. Alterations in the ACC in schizophrenia include perturbations in genes for vesicular glutamate transport and glutamate cycling [2, 10-12].

### Supplementary Methods

#### *Quality control demonstration of pyramidal neuron enrichment*

We have previously published extensive quality control studies for isolating pyramidal neuron population by LCM in human postmortem brains [11, 13-17]. The distinctive size and shape of pyramidal neurons allow for accurate capture of these cells by Nissl staining and morphological identification. We further performed quality control studies using qPCR and showed high levels of expression of neuronal markers neuron specific enolase (NSE) and vesicular glutamate transporter 1 (VGLUT1) in the enriched population of pyramidal neurons [13, 14, 16]. Our results demonstrated that highly enriched pyramidal neuron population can be captured following Nissl staining and morphological identification of cells.

#### *Pathway enrichment analysis*

The differential expression genes ranked based on statistics were used to map for the enrichment of different gene ontology term in Biological pathway (GOBP), Molecular Function (GOMF) and Cellular component (GOCC) categories using default GSEA parameters. An updated list of GO terms for all the three categories was downloaded from Bader lab (http://download.baderlab.org/EM_Genesets/). To investigate the effect of schizophrenia pathology for all comparisons, the normalized enrichment score of significant pathways (P-value<0.01) was used to generate heatmaps. Our list of GO terms for specific comparisons was large, so in order to reduce and catalog it into interpretable format we clustered the pathways based on biological themes. The theme for a given pathway was selected either based on text search where the name of the theme was used as keyword for text query, or based on the parent-child association between GO terms in our list of significant pathways (child pathways) and handpicked pathways in GO database (parent pathway) representing the theme, using GOdb package in R.

#### *Concordance score calculation in confirmation study*

To calculate concordance scores between datasets, a Pearson correlation analysis was conducted using the $\log_2$ fold change values. Using the list of genes of interest (56 genes), the values of $\log_2$ fold change were extracted from each dataset. For each correlation analysis between two datasets, the test was applied using only the observations of genes that were found in both datasets. The correlation coefficients of these are displayed as a graph that represents the correlation matrix for each dataset, where red represents negative correlation and blue represents positive correlation. Size and color intensity of the circles represent the level of similarity (Fig. 4).

***Parallel positive control for antipsychotic analysis***

In order to avoid high percentage matching among profiles treated with the same drugs, a consensus gene profile was generated by combining 8 clozapine treatment datasets (GSE48955: CLZ 3mg/kg 1h (duplicates), GSE48955: CLZ 3mg/kg 2h (duplicates), GSE48955: CLZ 8h, GSE33822: CLZ forebrain B6, GSE33822: CLZ hindbrain B6, GSE33822: CLZ wholebrain B6) based on p-values using meta-P package in R. The consensus gene profile was used to compare all 51 datasets to calculate the AUC scores. Adjusted P value lower than 0.05 is considered significant.

**Supplementary Results**

***Antipsychotic positive control analysis***

We further performed a positive control analysis for the antipsychotic treatment using a consensus gene profile by combining 8 clozapine treatment datasets with different doses, treatment lengths, and brain regions, as described in Supplementary Methods. The consensus gene list was compared with the 51 antipsychotic treatment profiles. We detected 36/51 upregulated and 21/51 downregulated antipsychotic with significant similarity to the consensus. In addition, 3/51 upregulated and 3/51 downregulated profiles showed AUC values higher than 0.85 and were associated with clozapine treatments (Fig. 3 blue curves). These suggest that the dataset was not influenced by the any antipsychotic treatment.

***Additional confirmation study***

Among all the datasets compared, we detected 15 genes that have the same directionality with at least 4 other data sets (upregulated: TNRC6C, LONRF1, HPGD, DNAH12, SERPINB5, SERTAD1, XTLT2, ABCF1 and PELI2; downregulated: BRI3BP, NPM1, B4GALT3, INF521, TMEM141, and PNKD) and 13 genes have the opposite directionality with at least 4 other datasets (upregulated: TBP, FRA10AC1, ABCF1, NRN1, and PARVA; downregulated: ZNF609, ODF4,

ZNF501, LPCAT3, LOXL2, B4GALT3, LSM14A, and TAS2R14). In the region and cell level combined comparisons, we detected 7 genes have the same directionality with at least 6 other data sets (upregulated: PELI2, TNRC6C, LONRF1, SERPINB5, and XYLT2; downregulated: NOX4 and RPS13) and 15 genes have the opposite directionality with at least 6 other data sets (upregulated: NRN1, TBP, ZBBX, ABCF1, FPA10AC1, and SERTAD1; downregulated: TAS2R14, TSPAN15, ZNF609, SLC11A1, LPCAT3, B4GALT3, LOXL2, LSM14A, and AP4B1) (Fig. 4).

Concordance scores were calculated in between our results and the other 16 region and cell level studies. We found out that the 3 comparisons of our data had the highest concordance scores. Consistent with our pathway analysis, we observed higher concordance in between deep neurons and the combined comparison, rather than superficial. Our results have positive concordance score with 4/11 datasets. 2/4 are iPSCs of neurons and progenitor neurons [18]. 1/4 is from a previously reported ACC dataset [19]. We also surprisingly detected positive concordance with a dataset from peripheral blood mononuclear cells. [20]. The datasets from DLPFC pyramidal neurons [3, 21] showed positive concordance with our deep neurons, but negative concordance with superficial neurons (Fig. 4).

**Supplementary discussion**

***Purity of pyramidal neuron population by LCM***

One may think that the large amount of immune related pathways are not directly from neurons, but is from microglia contamination, which is present low levels in LCM samples. However, neurons typically express chemokines and chemokine receptors [22]. For example, the regulation of neuronal chemokine fractalkine (CX3CL1) plays an important role for neuron survival [23]. Neuronal cytokine production has been studied for more than 2 decades [24], although it has not been comprehensively explored. In addition, we described our capacity to successfully purify pyramidal neurons in the supplementary methods [11, 13-17]. Therefore, our results demonstrated that cytokine and chemokine production was not exclusively from brain glial cells and neuronal cell derived cytokines and chemokines also play a role in this network in the brain. In our GSEA results, we observe 57% leukocyte chemotaxis pathways in all immunity related pathways, which indicates the dominant role of these pathways in neurons.

#### *Protein kinase deficits in schizophrenia*

Although protein kinases have been linked to schizophrenia pathophysiology for decades [25], the linkage between protein modification and schizophrenia pathophysiology at the transcriptional level has not been extensively explored. Mitogen-activated protein kinases were elevated in schizophrenia at the regional level [25], indicating dysregulated MAPK/ERK pathways in schizophrenia. In addition, our previous kinome analyses have also shown abnormal AKT activity in the ACC in schizophrenia [26]. A recent study showed that the dysregulated unfolded protein response observed in schizophrenia patients is mediated by endoplasmic reticulum transmembrane stress sensor pathways, which include protein kinase RNA-like ER kinase and p-JNK2 [27]. Taken together, these studies highlight the importance of kinases in schizophrenia pathophysiology.

#### *Concordance scores in confirmation study*

We compared our data with the previously reported DLPFC layer 3 and layer 5 pyramidal neuron profiles, as these datasets are the most similar to our experimental design. However, the concordance scores between these datasets and ours are not particularly high. While multiple reasons could have caused this observation, as we described at the beginning of the discussion, we think that the difference of gene expression profiles among different brain regions is the most reasonable explanation.
