## Supplementary Table 1 for "Transcriptional profile of pyramidal neurons in chronic schizophrenia reveals lamina-specific dysfunction of neuronal immunity"

**Supplementary table 1 Top differential expressed gene in the 3 indicated comparisons**

**Super_Deep upregulated**

| Gene name | logFC | AveExpr | P.Value |
|---|---|---|---|
| ANKRD20A1 | 0.771481 | 2.934472 | 1.45E-05 |
| MTND3P19 | 1.212009 | 3.480745 | 2.90E-05 |
| NRN1 | 0.717292 | 5.376916 | 0.000335 |
| PUM3 | 0.388281 | 3.101698 | 0.000715 |
| DMAC1 | 0.453668 | 4.600006 | 0.001065 |
| SERTAD1 | 0.343038 | 5.150986 | 0.001122 |
| RPL10AP2 | 0.399501 | 3.917066 | 0.001176 |
| LONRF1 | 0.39188 | 5.189656 | 0.001331 |
| ARL15 | 0.375465 | 3.335247 | 0.001402 |
| CCDC138 | 0.359536 | 2.909713 | 0.001489 |
| PARVA | 0.392535 | 4.459493 | 0.001801 |
| MTND4LP10 | 0.464169 | 2.15814 | 0.001802 |
| PELI2 | 0.342934 | 5.678829 | 0.002044 |
| MTATP6P17 | 0.413899 | 2.22237 | 0.002184 |
| THAP5 | 0.697433 | 4.596419 | 0.002372 |
| SERPINB5 | 0.376054 | 2.95327 | 0.002417 |
| POTEF | 0.851096 | 2.638289 | 0.002417 |
| PSG11 | 0.510747 | 3.247184 | 0.002573 |
| AC091868.1 | 0.442131 | 2.993311 | 0.002708 |
| TBP | 0.334164 | 4.946911 | 0.002984 |
| FRA10AC1 | 0.47514 | 3.364267 | 0.003029 |
| ZBBX | 0.366563 | 3.669518 | 0.003144 |
| IFNA6 | 0.485798 | 4.18425 | 0.00323 |
| XYLT2 | 0.335638 | 4.552123 | 0.003255 |
| HPGD | 0.351351 | 3.036439 | 0.003394 |
| ZNHIT2 | 0.50823 | 4.984113 | 0.003934 |
| UBE2MP1 | 0.730867 | 3.69478 | 0.003954 |
| CLIC3 | 0.466601 | 4.163422 | 0.004097 |
| TNRC6C | 0.285301 | 4.407275 | 0.004132 |
| ABCF1 | 0.895283 | 2.532438 | 0.004281 |
| DNAH12 | 0.333903 | 3.044927 | 0.004328 |
| TBCA | 0.653204 | 8.223674 | 0.004445 |
| AL109766.1 | 1.381995 | 6.944693 | 0.004462 |
| C11orf58 | 0.512052 | 2.177447 | 0.00463 |
| DDX5 | 1.032492 | 7.342541 | 0.004655 |
| GCAT | 0.308707 | 6.389977 | 0.004698 |
| KRAS | 0.430018 | 4.242529 | 0.004715 |
| TTBK2 | 0.276926 | 5.178771 | 0.00488 |
| EGLN2 | 0.250517 | 5.28797 | 0.004882 |
| MTND6P15 | 0.690543 | 2.564133 | 0.005007 |
| SAA2 | 0.78875 | 4.074 | 0.005179 |
| RFTN2 | 0.323868 | 3.882463 | 0.005415 |
| POLR2B | 0.501424 | 4.58082 | 0.005422 |
| TPRN | 0.292898 | 5.036924 | 0.005441 |

**Super_Deep downregulated**

| Gene name | logFC | AveExpr | P.Value |
|---|---|---|---|
| COLEC10 | -0.37834 | 3.719783 | 0.000158 |
| NPM1 | -0.83117 | 3.844913 | 0.000231 |
| YBX1P2 | -0.47959 | 4.635862 | 0.000349 |
| OR7E35P | -0.54104 | 5.237738 | 0.000479 |
| BRI3BP | -0.36846 | 5.203522 | 0.000501 |
| DDX39B | -0.97463 | 4.223606 | 0.000588 |
| DRD5P1 | -0.37696 | 5.087505 | 0.000715 |
| CD28 | -0.53943 | 4.134102 | 0.000732 |
| OR5BS1P | -0.49266 | 4.480863 | 0.000826 |
| AC087203.3 | -0.90366 | 4.870688 | 0.000971 |
| CNN1 | -0.43068 | 6.02295 | 0.001018 |
| SLC11A1 | -0.4283 | 5.023796 | 0.001026 |
| TSPAN15 | -0.29289 | 5.890792 | 0.001254 |
| GOLGA6L6 | -3.33014 | 5.332998 | 0.001275 |
| TMEM141 | -0.32322 | 5.991977 | 0.001374 |
| MS4A8 | -0.43653 | 5.075835 | 0.001481 |
| LSM14A | -0.58418 | 6.263129 | 0.00152 |
| LPCAT3 | -0.42744 | 4.735183 | 0.001755 |
| LINC02145 | -0.39878 | 4.299546 | 0.001783 |
| RPS13 | -1.87321 | 6.024963 | 0.001906 |
| ZNF521 | -0.40933 | 4.813524 | 0.001925 |
| C2 | -0.47184 | 5.089597 | 0.001983 |
| ZNF501 | -0.4831 | 4.816552 | 0.002099 |
| B4GALT3 | -0.30592 | 4.895337 | 0.002245 |
| AP4B1 | -0.40392 | 4.141355 | 0.002261 |
| NOX4 | -0.35514 | 4.551851 | 0.002356 |
| PNKD | -0.36009 | 4.679785 | 0.002372 |
| OR6T1 | -0.41358 | 5.105134 | 0.00263 |
| ODF4 | -0.31075 | 5.141804 | 0.002639 |
| CCL16 | -0.36546 | 4.115728 | 0.002674 |
| KIR3DL3 | -0.5983 | 5.760414 | 0.002753 |
| CD1C | -0.40584 | 5.163742 | 0.003064 |
| ACSM2A | -0.57939 | 4.803798 | 0.003142 |
| SPRR1B | -0.35817 | 6.815008 | 0.003229 |
| MBD3L2B | -0.5119 | 5.804773 | 0.003352 |
| TAS2R14 | -0.35213 | 4.136662 | 0.003398 |
| DKKL1 | -0.4171 | 4.836442 | 0.003445 |
| LOXL2 | -0.34139 | 5.050402 | 0.003456 |
| ZNF585B | -0.4556 | 6.160155 | 0.004013 |
| DDX11 | -0.28255 | 5.578598 | 0.004136 |
| ZNF609 | -0.33462 | 4.87469 | 0.004193 |
| SUFU | -0.24578 | 4.283835 | 0.004216 |
| ALOXE3 | -0.30082 | 5.464037 | 0.004348 |
| WDR75 | -0.35326 | 3.298658 | 0.004423 |

| | | | |
|---|---|---|---|
| TBC1D12 | 0.307337 | 3.598072 | 0.005539 |
| NEMP2 | 0.31415 | 2.615671 | 0.005573 |
| MTND6P2 | 0.29243 | 2.117628 | 0.005586 |
| MTND1P22 | 0.788805 | 3.325084 | 0.005739 |
| RDH14 | 0.506327 | 4.36872 | 0.005766 |
| KRT7 | 0.433653 | 5.251423 | 0.005992 |

**Superficial upregulated**

| Gene name | logFC | AveExpr | P.Value |
|---|---|---|---|
| ANKRD20A1 | 0.403774 | 2.934472 | 0.000576 |
| MTND3P19 | 0.625267 | 3.480745 | 0.001081 |
| NRN1 | 0.369964 | 5.376916 | 0.005537 |
| PUM3 | 0.199812 | 3.101698 | 0.009244 |
| DMAC1 | 0.238233 | 4.600006 | 0.01045 |
| MTND4LP10 | 0.252594 | 2.15814 | 0.011776 |
| SERTAD1 | 0.177786 | 5.150986 | 0.011814 |
| RPL10AP2 | 0.206804 | 3.917066 | 0.012279 |
| CCDC138 | 0.188584 | 2.909713 | 0.01316 |
| LONRF1 | 0.200104 | 5.189656 | 0.014526 |
| MTATP6P17 | 0.22015 | 2.22237 | 0.015567 |
| ARL15 | 0.188991 | 3.335247 | 0.016413 |
| PELI2 | 0.179218 | 5.678829 | 0.016604 |
| AC091868.1 | 0.234967 | 2.993311 | 0.018093 |
| PSG11 | 0.269711 | 3.247184 | 0.01816 |
| SERPINB5 | 0.196632 | 2.95327 | 0.018479 |
| POTEF | 0.442639 | 2.638289 | 0.019079 |
| PARVA | 0.197556 | 4.459493 | 0.019195 |
| TBP | 0.175941 | 4.946911 | 0.020396 |
| ZBBX | 0.193982 | 3.669518 | 0.020508 |
| THAP5 | 0.353249 | 4.596419 | 0.021968 |
| HPGD | 0.184466 | 3.036439 | 0.022574 |
| FRA10AC1 | 0.246115 | 3.364267 | 0.022609 |
| XYLT2 | 0.174993 | 4.552123 | 0.022839 |
| CLIC3 | 0.246867 | 4.163422 | 0.024501 |
| EGLN2 | 0.134639 | 5.28797 | 0.025308 |
| C11orf58 | 0.268682 | 2.177447 | 0.027783 |
| MTND6P15 | 0.36332 | 2.564133 | 0.028841 |
| TTBK2 | 0.144862 | 5.178771 | 0.029227 |
| TNRC6C | 0.145974 | 4.407275 | 0.029481 |
| ABCF1 | 0.459443 | 2.532438 | 0.029683 |
| AL109766.1 | 0.709847 | 6.944693 | 0.030334 |
| GCAT | 0.158968 | 6.389977 | 0.030939 |
| KRAS | 0.221102 | 4.242529 | 0.031252 |
| IFNA6 | 0.238468 | 4.18425 | 0.031394 |
| DNAH12 | 0.169648 | 3.044927 | 0.03147 |
| NEMP2 | 0.164659 | 2.615671 | 0.03156 |

| | | | |
|---|---|---|---|
| TXLNA | -0.32922 | 5.288131 | 0.004467 |
| CYP11A1 | -0.36718 | 4.534895 | 0.004709 |
| MTND4P28 | -0.5984 | 4.896494 | 0.004725 |
| PPIAP70 | -0.3356 | 3.231823 | 0.004746 |
| PIP5K1P1 | -0.40142 | 4.4156 | 0.00478 |
| AC090099.1 | -0.31612 | 4.640613 | 0.005002 |

**Superficial downregulated**

| Gene name | logFC | AveExpr | P.Value |
|---|---|---|---|
| NPM1 | -0.43524 | 3.844913 | 0.003826 |
| COLEC10 | -0.19118 | 3.719783 | 0.004008 |
| YBX1P2 | -0.25693 | 4.635862 | 0.004207 |
| BRI3BP | -0.19244 | 5.203522 | 0.006567 |
| OR7E35P | -0.27854 | 5.237738 | 0.007099 |
| DDX39B | -0.49532 | 4.223606 | 0.008902 |
| CD28 | -0.27625 | 4.134102 | 0.009715 |
| OR5BS1P | -0.25305 | 4.480863 | 0.010284 |
| CNN1 | -0.22331 | 6.02295 | 0.011054 |
| AC087203.3 | -0.46555 | 4.870688 | 0.011194 |
| SLC11A1 | -0.22166 | 5.023796 | 0.011249 |
| DRD5P1 | -0.18446 | 5.087505 | 0.01299 |
| LSM14A | -0.30386 | 6.263129 | 0.014076 |
| TMEM141 | -0.16613 | 5.991977 | 0.014221 |
| PNKD | -0.19523 | 4.679785 | 0.014616 |
| TSPAN15 | -0.147 | 5.890792 | 0.01558 |
| RPS13 | -0.97731 | 6.024963 | 0.016023 |
| C2 | -0.24705 | 5.089597 | 0.016091 |
| MS4A8 | -0.21974 | 5.075835 | 0.016972 |
| LPCAT3 | -0.21897 | 4.735183 | 0.016985 |
| ZNF521 | -0.21137 | 4.813524 | 0.017174 |
| GOLGA6L6 | -1.63696 | 5.332998 | 0.01788 |
| ZNF501 | -0.24958 | 4.816552 | 0.018107 |
| AP4B1 | -0.21036 | 4.141355 | 0.018115 |
| LINC02145 | -0.20076 | 4.299546 | 0.019039 |
| B4GALT3 | -0.15644 | 4.895337 | 0.020076 |
| ODF4 | -0.16134 | 5.141804 | 0.020402 |
| SPRR1B | -0.18972 | 6.815008 | 0.020758 |
| KIR3DL3 | -0.31081 | 5.760414 | 0.020904 |
| CD1C | -0.21181 | 5.163742 | 0.021825 |
| NOX4 | -0.17966 | 4.551851 | 0.022026 |
| OR6T1 | -0.211 | 5.105134 | 0.022496 |
| ACSM2A | -0.30103 | 4.803798 | 0.022759 |
| MBD3L2B | -0.26692 | 5.804773 | 0.023267 |
| TAS2R14 | -0.18255 | 4.136662 | 0.024242 |
| TXLNA | -0.17561 | 5.288131 | 0.024825 |
| DDX11 | -0.14855 | 5.578598 | 0.025525 |

| | | | | | | | |
|---|---|---|---|---|---|---|---|
| UBE2MP1 | 0.366104 | 3.69478 | 0.031988 | LINC01561 | -0.18035 | 5.343888 | 0.025902 |
| MTND6P2 | 0.1526 | 2.117628 | 0.032325 | DKKL1 | -0.21389 | 4.836442 | 0.025951 |
| COX6B1P3 | 0.16504 | 4.514565 | 0.032768 | CCL16 | -0.18091 | 4.115728 | 0.026833 |
| ZNHIT2 | 0.252805 | 4.984113 | 0.033043 | SUFU | -0.12803 | 4.283835 | 0.02717 |
| DDX5 | 0.523309 | 7.342541 | 0.033343 | MCM5 | -0.13706 | 5.234862 | 0.027678 |
| CCDC77 | 0.626366 | 7.718285 | 0.033566 | MTND4P28 | -0.31461 | 4.896494 | 0.027866 |
| TBC1D12 | 0.158293 | 3.598072 | 0.034332 | ZNF609 | -0.17258 | 4.87469 | 0.028535 |
| KRT10 | 0.187571 | 4.630333 | 0.034547 | ZNF585B | -0.23332 | 6.160155 | 0.028801 |
| RDH14 | 0.261328 | 4.36872 | 0.034863 | LOXL2 | -0.17006 | 5.050402 | 0.030325 |
| RFTN2 | 0.165544 | 3.882463 | 0.035143 | FAM170A | -0.18089 | 4.548002 | 0.031593 |
| AC025627.4 | 0.454547 | 5.258369 | 0.035314 | C20orf78 | -0.20785 | 4.332229 | 0.031843 |
| ORC2 | 0.158456 | 3.431943 | 0.035717 | SREBF2 | -0.22235 | 4.583029 | 0.031979 |
| MTND1P22 | 0.404813 | 3.325084 | 0.035749 | ALOXE3 | -0.15218 | 5.464037 | 0.032261 |

**Deep upregulated**

| Gene name | logFC | AveExpr | P.Value |
|---|---|---|---|
| ANKRD20A1 | 0.367706 | -2.93447 | 0.001094 |
| MTND3P19 | 0.586742 | -3.48074 | 0.001464 |
| NRN1 | 0.347328 | -5.37692 | 0.006953 |
| PUM3 | 0.188469 | -3.1017 | 0.010993 |
| ARL15 | 0.186473 | -3.33525 | 0.014505 |
| SERTAD1 | 0.165251 | -5.15099 | 0.015269 |
| LONRF1 | 0.191776 | -5.18966 | 0.015392 |
| RPL10AP2 | 0.192698 | -3.91707 | 0.015593 |
| DMAC1 | 0.215435 | -4.60001 | 0.016169 |
| PARVA | 0.194979 | -4.45949 | 0.017018 |
| CCDC138 | 0.170952 | -2.90971 | 0.019622 |
| THAP5 | 0.344183 | -4.59642 | 0.021075 |
| IFNA6 | 0.247329 | -4.18425 | 0.021534 |
| PELI2 | 0.163716 | -5.67883 | 0.023197 |
| POTEF | 0.408456 | -2.63829 | 0.024934 |
| POLR2B | 0.263458 | -4.58082 | 0.025672 |
| SERPINB5 | 0.179422 | -2.95327 | 0.025739 |
| ZNHIT2 | 0.255425 | -4.98411 | 0.026332 |
| MTATP6P17 | 0.193749 | -2.22237 | 0.026864 |
| UBE2MP1 | 0.364763 | -3.69478 | 0.027421 |
| MTND4LP10 | 0.211575 | -2.15814 | 0.027739 |
| TBCA | 0.329855 | -8.22367 | 0.027791 |
| FRA10AC1 | 0.229025 | -3.36427 | 0.02793 |
| PSG11 | 0.241036 | -3.24718 | 0.028324 |
| XYLT2 | 0.160645 | -4.55212 | 0.030271 |
| AC091868.1 | 0.207164 | -2.99331 | 0.030299 |
| TBP | 0.158223 | -4.94691 | 0.03039 |
| DNAH12 | 0.164255 | -3.04493 | 0.031316 |
| TNRC6C | 0.139327 | -4.40727 | 0.031581 |
| ZBBX | 0.172581 | -3.66952 | 0.032265 |

**Deep downregulated**

| Gene name | logFC | AveExpr | P.Value |
|---|---|---|---|
| COLEC10 | -0.18716 | -3.71978 | 0.003624 |
| NPM1 | -0.39593 | -3.84491 | 0.006244 |
| DRD5P1 | -0.1925 | -5.0875 | 0.007703 |
| OR7E35P | -0.26251 | -5.23774 | 0.008577 |
| DDX39B | -0.47931 | -4.22361 | 0.008877 |
| YBX1P2 | -0.22266 | -4.63586 | 0.009659 |
| BRI3BP | -0.17602 | -5.20352 | 0.009836 |
| CD28 | -0.26319 | -4.1341 | 0.010781 |
| GOLGA6L6 | -1.69318 | -5.333 | 0.011706 |
| OR5BS1P | -0.23961 | -4.48086 | 0.011873 |
| TSPAN15 | -0.14588 | -5.89079 | 0.013244 |
| AC087203.3 | -0.43811 | -4.87069 | 0.013414 |
| SLC11A1 | -0.20664 | -5.0238 | 0.014312 |
| CNN1 | -0.20737 | -6.02295 | 0.014427 |
| MS4A8 | -0.21679 | -5.07584 | 0.015028 |
| TMEM141 | -0.15709 | -5.99198 | 0.016387 |
| LINC02145 | -0.19802 | -4.29955 | 0.01694 |
| LPCAT3 | -0.20846 | -4.73518 | 0.018687 |
| LSM14A | -0.28032 | -6.26313 | 0.018827 |
| CCL16 | -0.18455 | -4.11573 | 0.019934 |
| ZNF521 | -0.19795 | -4.81352 | 0.020787 |
| NOX4 | -0.17548 | -4.55185 | 0.020838 |
| B4GALT3 | -0.14948 | -4.89534 | 0.021548 |
| ZNF501 | -0.23351 | -4.81655 | 0.02198 |
| RPS13 | -0.8959 | -6.02496 | 0.022014 |
| C2 | -0.2248 | -5.0896 | 0.023044 |
| OR6T1 | -0.20257 | -5.10513 | 0.02346 |
| AP4B1 | -0.19356 | -4.14135 | 0.024136 |
| LOXL2 | -0.17134 | -5.0504 | 0.024397 |
| ODF4 | -0.14941 | -5.1418 | 0.02603 |

| | | | | | | | |
|---|---|---|---|---|---|---|---|
| HPGD | 0.166885 | -3.03644 | 0.03228 | KIR3DL3 | -0.28749 | -5.76041 | 0.026803 |
| DDX5 | 0.509183 | -7.34254 | 0.032365 | DKKL1 | -0.2032 | -4.83644 | 0.028568 |
| ABCF1 | 0.43584 | -2.53244 | 0.032797 | ACSM2A | -0.27836 | -4.8038 | 0.029056 |
| GTF2A1 | 0.161686 | -5.3642 | 0.033608 | CD1C | -0.19403 | -5.16374 | 0.029356 |
| AL109766.1 | 0.672148 | -6.94469 | 0.033803 | WDR75 | -0.1762 | -3.29866 | 0.029558 |
| SAA2 | 0.390362 | -4.074 | 0.034004 | PPIAP70 | -0.16878 | -3.23182 | 0.029628 |
| KRT7 | 0.217457 | -5.25142 | 0.034958 | TAS2R14 | -0.16958 | -4.13666 | 0.03009 |
| KRAS | 0.208916 | -4.24253 | 0.035158 | ALOXE3 | -0.14864 | -5.46404 | 0.0307 |
| GCAT | 0.149739 | -6.38998 | 0.03536 | MBD3L2B | -0.24498 | -5.80477 | 0.030831 |
| TPRN | 0.144146 | -5.03692 | 0.036062 | ZNF585B | -0.22228 | -6.16016 | 0.031163 |
| KMT2C | 0.111089 | -4.57568 | 0.036379 | PIP5K1P1 | -0.19959 | -4.4156 | 0.031565 |
| TRPM5 | 0.150086 | -4.58654 | 0.036387 | PNKD | -0.16486 | -4.67978 | 0.031636 |
| RFTN2 | 0.158325 | -3.88246 | 0.037144 | OR8B8 | -0.25534 | -5.14043 | 0.032006 |
| RGS6 | 0.183165 | -4.43785 | 0.03746 | CYP11A1 | -0.18112 | -4.53489 | 0.032557 |
| CLIC3 | 0.219734 | -4.16342 | 0.037629 | DEFB110 | -0.14167 | -3.33491 | 0.032658 |
| SNORD114-2 | 0.555403 | -3.71292 | 0.038086 | SIRPB2 | -0.14731 | -5.19113 | 0.032717 |
| UCK1 | 0.244888 | -4.69841 | 0.038595 | SPRR1B | -0.16844 | -6.81501 | 0.032951 |
| C11orf58 | 0.24337 | -2.17745 | 0.038688 | ZNF609 | -0.16204 | -4.87469 | 0.033253 |
| TBC1D12 | 0.149044 | -3.59807 | 0.039145 | TP53AIP1 | -0.22018 | -4.19418 | 0.033641 |
| TTBK2 | 0.132064 | -5.17877 | 0.039277 | ACCSL | -0.14104 | -5.58814 | 0.034359 |
