## Supplementary Table 2 for "Transcriptional profile of pyramidal neurons in chronic schizophrenia reveals lamina-specific dysfunction of neuronal immunity"

### Supplementary Table 2 Pathways involved in schizophrenia pathophysiology through GSEA

| Pathway | SCZvsCtrl | SCZvsCtrl_D | SCZvsCtrl_S |
|---|---|---|---|
| **Transcription** | | | |
| POSTTRANSCRIPTIONAL REGULATION OF GENE EXPRESSION | 0 | 0.2971033 | 0 |
| REGULATION OF DNA-TEMPLATED TRANSCRIPTION IN RESPONSE TO STRESS | 0.4209953 | 0.4128795 | 0.3492673 |
| REGULATION OF RNA POLYMERASE II PROMOTER IN RESPONSE TO HYPOXIA | 0.4656593 | 0.45597392 | 0.37566537 |
| REGULATION OF RNA POLYMERASE II PROMOTER IN RESPONSE TO STRESS | 0.4350208 | 0.42602116 | 0 |
| REGULATION OF VIRAL TRANSCRIPTION | 0.4513928 | 0.45089537 | 0 |
| REPRESSING TRANSCRIPTION FACTOR BINDING | 0.42931 | 0.4179329 | 0 |
| RNA POLYMERASE II REPRESSING TRANSCRIPTION FACTOR BINDING | 0.5278212 | 0 | 0.5335609 |
| TRANSCRIPTION-COUPLED NUCLEOTIDE-EXCISION REPAIR | 0 | 0.41460714 | 0 |
| VIRAL TRANSCRIPTION | 0 | 0 | -0.41784468 |
| **Translation** | | | |
| COTRANSLATIONAL PROTEIN TARGETING TO MEMBRANE | 0.4228422 | 0.43597224 | -0.41340497 |
| CYTOPLASMIC TRANSLATION | 0.4916354 | 0.51095414 | 0 |
| CYTOPLASMIC TRANSLATIONAL INITIATION | 0.6206517 | 0 | 0 |
| MITOCHONDRIAL TRANSLATION | 0.3910002 | 0.39043027 | 0.41419154 |
| MITOCHONDRIAL TRANSLATIONAL ELONGATION | 0 | 0 | 0.40039274 |
| MITOCHONDRIAL TRANSLATIONAL TERMINATION | 0 | 0.41739705 | 0.4173968 |
| SRP-DEPENDENT COTRANSLATIONAL PROTEIN TARGETING TO MEMBRANE | 0.4354171 | 0.4511298 | -0.4253419 |
| TRANSLATION FACTOR ACTIVITY, RNA BINDING | 0.448428 | 0.44332832 | 0 |
| TRANSLATION | 0.4017877 | 0.4067169 | 0 |
| TRANSLATIONAL ELONGATION | 0.3896833 | 0.40231618 | 0.38191894 |
| TRANSLATIONAL INITIATION | 0.4820776 | 0.48899695 | 0 |
| TRANSLATIONAL TERMINATION | 0.4348054 | 0.4294353 | 0.43336582 |
| **Protein Modification** | | | |
| PEPTIDYL-THREONINE PHOSPHORYLATION | 0 | 0 | 0.4143322 |
| POSITIVE REGULATION OF ERK1 AND ERK2 CASCADE | 0 | 0 | -0.33259064 |
| PROTEIN K11-LINKED UBIQUITINATION | 0 | 0.5870458 | 0 |
| PROTEIN OXIDATION | 0 | 0 | -0.67212313 |
| PROTEIN POLYUBIQUITINATION | 0.3529768 | 0.3593769 | 0.3106512 |
| REGULATION OF ERK1 AND ERK2 CASCADE | -0.3017069 | -0.2856738 | 0 |
| REGULATION OF PROTEIN FOLDING | 0 | 0 | 0.74522674 |
| **ATP/Respi** | | | |
| ATP METABOLIC PROCESS | 0 | 0.35959795 | 0 |
| ATPASE ACTIVITY | 0.295576 | 0.2924602 | 0 |
| CELLULAR RESPIRATION | 0.3568024 | 0 | 0.33658525 |
| RESPIRATORY BURST | 0 | 0 | -0.666244 |
| **Mitochondria** | | | |
| INNER MITOCHONDRIAL MEMBRANE PROTEIN COMPLEX | 0.3965123 | 0 | 0 |
| INTEGRAL COMPONENT OF MITOCHONDRIAL MEMBRANE | 0.4577092 | 0.4601709 | 0.43534383 |
| INTRINSIC COMPONENT OF MITOCHONDRIAL MEMBRANE | 0.4346886 | 0.4374556 | 0 |
| MITOCHONDRIAL GENE EXPRESSION | 0 | 0 | 0.35573474 |
| MITOCHONDRIAL INNER MEMBRANE | 0.2938931 | 0.29766372 | 0.2661932 |
| MITOCHONDRIAL MEMBRANE PART | 0.4101473 | 0.4055913 | 0.37418398 |
| MITOCHONDRIAL OUTER MEMBRANE | 0 | 0 | 0 |
| MITOCHONDRIAL PROTEIN COMPLEX | 0.3826416 | 0.37726352 | 0.35840493 |
| MITOCHONDRIAL RIBOSOME | 0 | 0 | 0.3965855 |
| OUTER MITOCHONDRIAL MEMBRANE PROTEIN COMPLEX | 0 | 0.7053745 | 0.7071472 |
| **ER** | | | |
| ESTABLISHMENT OF PROTEIN LOCALIZATION TO ENDOPLASMIC RETICULUM | 0.427108 | 0.44145682 | 0 |
| PROTEIN LOCALIZATION TO ENDOPLASMIC RETICULUM | 0.41424 | 0.42070618 | 0 |
| **Organelle** | | | |
| ESTABLISHMENT OF PROTEIN LOCALIZATION TO ORGANELLE | 0 | 0.29687834 | 0 |
| INTRINSIC COMPONENT OF ORGANELLE MEMBRANE | 0 | 0.33775064 | 0.30660394 |
| ORGANELLE INNER MEMBRANE | 0 | 0.29181665 | 0 |
| **Cytoskeleton** | | | |
| ACTIN CYTOSKELETON ORGANIZATION | 0 | 0.283129 | 0 |
| CYTOSKELETON-DEPENDENT INTRACELLULAR TRANSPORT | 0.3595407 | 0.3798889 | 0.33396918 |
| ESTABLISHMENT OR MAINTENANCE OF ACTIN CYTOSKELETON POLARITY | 0.718092 | 0.7188997 | 0 |
| ESTABLISHMENT OR MAINTENANCE OF CYTOSKELETON POLARITY | 0.6387135 | 0.63730526 | 0 |

| | | | |
|---|---|---|---|
| GAMMA-TUBULIN BINDING | 0.5942764 | 0.5905096 | 0.5974093 |
| REGULATION OF MITOTIC SPINDLE ASSEMBLY | 0.6235224 | 0.6320686 | 0 |
| **Axon** | | | |
| NEURON PROJECTION CYTOPLASM | 0 | 0.40513325 | 0 |
| NEURON PROJECTION EXTENSION | 0 | 0.43597773 | 0 |
| NEURON SPINE | 0 | 0.3525331 | 0 |
| **Dendrite** | | | |
| DENDRITIC SPINE | 0 | 0.3448264 | 0 |
| **Synapse** | | | |
| ASYMMETRIC SYNAPSE | 0.3345696 | 0.34774503 | 0 |
| GLUTAMATERGIC SYNAPSE | 0 | 0.29175848 | 0 |
| INTEGRAL COMPONENT OF POSTSYNAPTIC DENSITY MEMBRANE | 0.5057465 | 0.5126403 | 0 |
| INTEGRAL COMPONENT OF POSTSYNAPTIC SPECIALIZATION MEMBRANE | 0.4342942 | 0.44216675 | 0.39557064 |
| INTRINSIC COMPONENT OF POSTSYNAPTIC DENSITY MEMBRANE | 0.4905787 | 0.49732748 | 0.45578596 |
| INTRINSIC COMPONENT OF POSTSYNAPTIC SPECIALIZATION MEMBRANE | 0 | 0.43284112 | 0 |
| NEURON TO NEURON SYNAPSE | 0.3318627 | 0.34773207 | 0 |
| POSTSYNAPTIC DENSITY MEMBRANE | 0.4595476 | 0.4659215 | 0.43943357 |
| POSTSYNAPTIC DENSITY | 0.3326058 | 0.34773958 | 0 |
| POSTSYNAPTIC MEMBRANE | 0.339933 | 0.33605674 | 0.3083371 |
| POSTSYNAPTIC SPECIALIZATION MEMBRANE | 0.4232878 | 0.42073014 | 0.38832527 |
| POSTSYNAPTIC SPECIALIZATION | 0.3266305 | 0.33731592 | 0 |
| SYNAPTIC MEMBRANE | 0 | 0.29524243 | 0.26660293 |
| **Receptor** | | | |
| C-C CHEMOKINE RECEPTOR ACTIVITY | -0.6164724 | -0.6206737 | 0 |
| CARGO RECEPTOR ACTIVITY | -0.4602123 | -0.46106815 | -0.43293548 |
| CCR CHEMOKINE RECEPTOR BINDING | 0 | 0 | -0.55911744 |
| CHEMOKINE RECEPTOR ACTIVITY | -0.6487981 | -0.6520509 | -0.58907795 |
| CYTOKINE RECEPTOR ACTIVITY | -0.4484277 | -0.46709996 | -0.4108187 |
| FIBROBLAST GROWTH FACTOR RECEPTOR SIGNALING PATHWAY | 0.4058012 | 0 | 0 |
| G PROTEIN-COUPLED CHEMOATTRACTANT RECEPTOR ACTIVITY | -0.6487981 | -0.6520509 | -0.58907795 |
| GABA RECEPTOR COMPLEX | 0 | 0 | 0.61050135 |
| ICOSANOID RECEPTOR ACTIVITY | 0.7225263 | 0.7181881 | 0.70172364 |
| POSITIVE REGULATION OF T CELL RECEPTOR SIGNALING PATHWAY | -0.6791645 | -0.6455166 | 0 |
| PROSTAGLANDIN RECEPTOR ACTIVITY | 0.7744611 | 0.75488657 | 0.7765301 |
| PROSTANOID RECEPTOR ACTIVITY | 0.7526635 | 0.7448913 | 0.7390884 |
| **Trasmission** | | | |
| REGULATION OF TRANSMISSION OF NERVE IMPULSE | -0.6769567 | -0.66421056 | -0.64967984 |
| **Receptor** | | | |
| SCAVENGER RECEPTOR ACTIVITY | -0.5563271 | -0.55456376 | -0.5208628 |
| SEROTONIN RECEPTOR SIGNALING PATHWAY | 0 | -0.48958847 | 0 |
| **Transporter** | | | |
| ACTIVE TRANSMEMBRANE TRANSPORTER ACTIVITY | -0.2891749 | -0.2867565 | 0 |
| AMINO ACID TRANSMEMBRANE TRANSPORTER ACTIVITY | -0.4345679 | -0.41611758 | -0.4265707 |
| AMMONIUM TRANSMEMBRANE TRANSPORTER ACTIVITY | -0.5443257 | -0.5454719 | 0 |
| ANION TRANSMEMBRANE TRANSPORT | -0.3051553 | -0.30251035 | 0 |
| ANION TRANSMEMBRANE TRANSPORTER ACTIVITY | -0.2939813 | -0.28741428 | 0 |
| CARBOXYLIC ACID TRANSMEMBRANE TRANSPORT | -0.3809365 | -0.37149093 | -0.3698196 |
| CARBOXYLIC ACID TRANSMEMBRANE TRANSPORTER ACTIVITY | -0.4207446 | -0.41139147 | -0.39691144 |
| CARBOXYLIC ACID TRANSPORT | -0.3171873 | -0.31038103 | -0.30298162 |
| CHOLESTEROL TRANSPORTER ACTIVITY | 0 | 0 | -0.6391872 |
| DICARBOXYLIC ACID TRANSMEMBRANE TRANSPORTER ACTIVITY | -0.5136235 | -0.50452155 | 0 |
| ICOSANOID TRANSPORT | 0 | -0.5237443 | 0 |
| L-AMINO ACID TRANSPORT | -0.4681799 | -0.44185326 | -0.46396026 |
| LIPID TRANSPORTER ACTIVITY | 0 | 0 | -0.3867698 |
| METAL ION TRANSMEMBRANE TRANSPORTER ACTIVITY | -0.2693176 | 0 | 0 |
| MONOCARBOXYLIC ACID TRANSMEMBRANE TRANSPORTER ACTIVITY | -0.4626368 | -0.4570776 | 0 |
| NEUROTRANSMITTER TRANSPORTER ACTIVITY | -0.4661535 | 0 | 0 |
| ORGANIC ACID TRANSMEMBRANE TRANSPORTER ACTIVITY | -0.4207446 | -0.41139147 | -0.39691144 |
| ORGANIC ANION TRANSMEMBRANE TRANSPORTER ACTIVITY | -0.3743136 | -0.35869086 | -0.3422472 |
| ORGANIC ANION TRANSPORT | -0.3097518 | -0.29605988 | -0.31191695 |
| ORGANIC CATION TRANSMEMBRANE TRANSPORTER ACTIVITY | 0 | -0.55816746 | 0 |

| | | | |
|---|---|---|---|
| ORGANIC HYDROXY COMPOUND TRANSMEMBRANE TRANSPORTER ACTIVITY | 0 | -0.43071973 | 0 |
| SECONDARY ACTIVE TRANSMEMBRANE TRANSPORTER ACTIVITY | -0.4119955 | -0.40480638 | -0.39796755 |
| SODIUM ION TRANSMEMBRANE TRANSPORTER ACTIVITY | -0.3250429 | 0 | 0 |
| STEROL TRANSPORTER ACTIVITY | 0 | 0 | -0.51992875 |
| SULFUR COMPOUND TRANSMEMBRANE TRANSPORTER ACTIVITY | -0.5925434 | -0.57877344 | -0.54759955 |
| **neurotransmitter** | | | |
| NEUROTRANSMITTER:SODIUM SYMPORTER ACTIVITY | -0.5734214 | -0.5630056 | 0 |
| **Secretion** | | | |
| ARACHIDONIC ACID SECRETION | 0 | -0.5406368 | 0 |
| ICOSANOID SECRETION | 0 | -0.5228797 | 0 |
| **Cytokine** | | | |
| NEGATIVE REGULATION OF CYTOKINE SECRETION | -0.4204528 | -0.40874472 | -0.4216135 |
| REGULATION OF CYTOKINE SECRETION | 0 | -0.3155167 | 0 |
| REGULATION OF INTERLEUKIN-8 SECRETION | -0.5804965 | -0.5851303 | 0 |
| CYTOKINE RECEPTOR ACTIVITY | -0.4484277 | -0.46709996 | -0.4108187 |
| CYTOKINE ACTIVITY | 0 | -0.3044843 | 0 |
| CYTOKINE BINDING | -0.3697091 | -0.38870886 | 0 |
| CYTOKINE PRODUCTION | -0.350553 | -0.34816307 | 0 |
| NEGATIVE REGULATION OF CYTOKINE PRODUCTION | -0.3364726 | -0.3198883 | -0.35391167 |
| NEGATIVE REGULATION OF INTERLEUKIN-8 PRODUCTION | 0 | -0.62104726 | 0 |
| NEGATIVE REGULATION OF TNF SUPERFAMILY CYTOKINE PRODUCTION | -0.4485381 | -0.43721634 | 0 |
| REGULATION OF INTERLEUKIN-10 PRODUCTION | -0.4658923 | -0.45012352 | -0.48708788 |
| REGULATION OF INTERLEUKIN-8 PRODUCTION | 0 | -0.42007706 | 0 |
| REGULATION OF MACROPHAGE CYTOKINE PRODUCTION | -0.7053993 | -0.6760612 | -0.6890065 |
| **Immune** | | | |
| LEUKOCYTE CHEMOTAXIS | -0.4218217 | -0.4234405 | -0.4171856 |
| LEUKOCYTE MIGRATION | -0.3006083 | -0.29767117 | 0 |
| LYMPHOCYTE CHEMOTAXIS | 0 | 0 | -0.5285425 |
| MONOCYTE CHEMOTAXIS | -0.5094023 | -0.49910438 | -0.56142104 |
| MONONUCLEAR CELL MIGRATION | -0.4896382 | -0.4799078 | -0.53843135 |
| MYELOID LEUKOCYTE MIGRATION | -0.3694409 | -0.36967045 | -0.37339365 |
| NEUTROPHIL CHEMOTAXIS | -0.3866253 | -0.39349398 | 0 |
| POSITIVE REGULATION OF LEUKOCYTE CHEMOTAXIS | -0.398333 | -0.38647282 | -0.4852239 |
| POSITIVE REGULATION OF LEUKOCYTE MIGRATION | 0 | 0 | -0.38535094 |
| POSITIVE REGULATION OF LYMPHOCYTE CHEMOTAXIS | 0 | 0 | -0.61327606 |
| POSITIVE REGULATION OF LYMPHOCYTE MIGRATION | 0 | 0 | -0.52468246 |
| POSITIVE REGULATION OF MONOCYTE CHEMOTAXIS | -0.6218818 | -0.61778307 | 0 |
| REGULATION OF LEUKOCYTE CHEMOTAXIS | 0 | -0.35291773 | -0.41133818 |
| REGULATION OF LYMPHOCYTE CHEMOTAXIS | 0 | 0 | -0.5771987 |
| REGULATION OF NEUTROPHIL MIGRATION | 0 | 0 | -0.49605244 |
| REGULATION OF T CELL CHEMOTAXIS | 0 | 0 | -0.6250676 |
| EOSINOPHIL CHEMOTAXIS | -0.7083336 | -0.7004597 | -0.70811224 |
| EOSINOPHIL MIGRATION | -0.697153 | -0.6875337 | -0.69835323 |
| GRANULOCYTE CHEMOTAXIS | -0.3965407 | -0.40334058 | 0 |
| GRANULOCYTE MIGRATION | -0.3791013 | 0 | -0.38834926 |
| C-C CHEMOKINE RECEPTOR ACTIVITY | -0.6164724 | -0.6206737 | 0 |
| CCR CHEMOKINE RECEPTOR BINDING | 0 | 0 | -0.55911744 |
| CHEMOKINE RECEPTOR ACTIVITY | -0.6487981 | -0.6520509 | -0.58907795 |
| ADAPTIVE IMMUNE RESPONSE | -0.3655469 | -0.35279164 | -0.37771082 |
| B CELL MEDIATED IMMUNITY | -0.4151166 | -0.39143702 | -0.43119422 |
| COMPLEMENT ACTIVATION, CLASSICAL PATHWAY | -0.5345565 | -0.53026235 | -0.5511242 |
| COMPLEMENT ACTIVATION | -0.4764351 | -0.47483796 | -0.51917 |
| REGULATION OF COMPLEMENT ACTIVATION | -0.4744149 | -0.46187505 | -0.54829776 |
| IMMUNOGLOBULIN BINDING | -0.6219181 | -0.63547 | -0.62739 |
| IMMUNOGLOBULIN MEDIATED IMMUNE RESPONSE | -0.4061147 | -0.39144573 | -0.42382815 |
| REGULATION OF HUMORAL IMMUNE RESPONSE | -0.428784 | -0.4263542 | -0.49670517 |
| REGULATION OF IMMUNE EFFECTOR PROCESS | -0.2801156 | -0.26785088 | -0.3001346 |
| HUMORAL IMMUNE RESPONSE MEDIATED BY CIRCULATING IMMUNOGLOBULIN | -0.5004132 | -0.4955407 | -0.5174437 |
| HUMORAL IMMUNE RESPONSE | -0.3426326 | -0.3455187 | -0.35544086 |
| NEGATIVE T CELL SELECTION | -0.7092772 | -0.6914009 | -0.6932712 |
| LYMPHOCYTE MEDIATED IMMUNITY | -0.3837055 | -0.36440825 | -0.4017144 |

| | | | |
|---|---|---|---|
| POSITIVE REGULATION OF T CELL PROLIFERATION | -0.3828556 | 0 | 0 |
| REGULATION OF LYMPHOCYTE PROLIFERATION | -0.3191193 | -0.3025959 | 0 |
| REGULATION OF T CELL PROLIFERATION | 0 | -0.3302253 | -0.34571287 |
| T CELL DIFFERENTIATION IN THYMUS | -0.4479866 | -0.43875417 | 0 |
| T CELL SELECTION | 0 | 0 | -0.50288785 |
| THYMIC T CELL SELECTION | -0.6284612 | -0.61653703 | 0 |
| ADAPTIVE IMMUNE RESPONSE BASED ON SOMATIC RECOMBINATION | -0.3799993 | -0.36083075 | -0.38979477 |
| POSITIVE REGULATION OF T CELL RECEPTOR SIGNALING PATHWAY | -0.6791645 | -0.6455166 | 0 |
| ANTIGEN PROCESSING AND PRESENTATION OF ENDOGENOUS ANTIGEN | 0 | 0 | -0.6458494 |
| ANTIGEN PROCESSING AND PRESENTATION OF EXOGENOUS ANTIGEN | 0.3567469 | 0.35665044 | 0 |
| ANTIGEN PROCESSING AND PRESENTATION OF EXOGENOUS PEPTIDE VIA MHC CLASS II | 0.4093595 | 0.39800212 | 0 |
| ANTIGEN PROCESSING AND PRESENTATION OF EXOGENOUS PEPTIDE ANTIGEN | 0.3854866 | 0.3820784 | 0 |
| ANTIGEN PROCESSING AND PRESENTATION OF PEPTIDE VIA MHC CLASS II | 0 | 0.39800212 | 0 |
| REGULATION OF HEMATOPOIETIC PROGENITOR CELL DIFFERENTIATION | 0.4678842 | 0.46775398 | 0.38886258 |
| REGULATION OF HEMATOPOIETIC STEM CELL DIFFERENTIATION | 0.5017693 | 0.5084241 | 0.41120362 |
| IMMUNE RESPONSE-ACTIVATING SIGNAL TRANSDUCTION | 0 | 0 | -0.2844257 |
| IMMUNE RESPONSE-REGULATING SIGNALING PATHWAY | 0 | 0 | -0.2763649 |
| ACTIVATION OF IMMUNE RESPONSE | 0 | -0.2567264 | -0.307885 |
| **Proteolysis** | | | |
| ANAPHASE-PROMOTING COMPLEX-DEPENDENT CATABOLIC PROCESS | 0.4805062 | 0.49522433 | 0 |
| MODIFICATION-DEPENDENT PROTEIN CATABOLIC PROCESS | 0.2840147 | 0.29391894 | 0 |
| PROTEASOMAL PROTEIN CATABOLIC PROCESS | 0.3052683 | 0.30749986 | 0 |
| PROTEASOMAL UBIQUITIN-INDEPENDENT PROTEIN CATABOLIC PROCESS | 0.6561729 | 0.6591135 | 0 |
| PROTEASOME-MEDIATED UBIQUITIN-DEPENDENT PROTEIN CATABOLIC PROCESS | 0 | 0.32165468 | 0 |
| PROTEIN DEUBIQUITINATION | 0 | 0.32102683 | 0 |
| PROTEIN MODIFICATION BY SMALL PROTEIN REMOVAL | 0 | 0.31958103 | 0 |
| UBIQUITIN-DEPENDENT PROTEIN CATABOLIC PROCESS | 0 | 0.2922979 | 0 |
| REGULATION OF PROTEIN PROCESSING | -0.4172971 | -0.40312278 | -0.4437227 |
| **Response to stress** | | | |
| CELLULAR RESPONSE TO HYPOXIA | 0.3533061 | 0.3493245 | 0 |
| DEFENSE RESPONSE TO BACTERIUM | -0.3983852 | -0.40737566 | -0.3664167 |
| DEFENSE RESPONSE TO GRAM-NEGATIVE BACTERIUM | -0.4740552 | -0.4753186 | -0.4377799 |
| DEFENSE RESPONSE TO GRAM-POSITIVE BACTERIUM | 0 | -0.38255495 | 0 |
| DEFENSE RESPONSE TO OTHER ORGANISM | -0.3290343 | -0.329963 | -0.30491275 |
| INFLAMMATORY RESPONSE | -0.2960659 | -0.29288608 | -0.293434 |
| REGULATION OF ACUTE INFLAMMATORY RESPONSE | -0.3789535 | -0.3725005 | -0.45540485 |
| REGULATION OF BLOOD COAGULATION | -0.3954463 | 0 | 0 |
| REGULATION OF INFLAMMATORY RESPONSE | 0 | 0 | -0.3062746 |
| RESPONSE TO INTERFERON-GAMMA | 0 | 0 | -0.36389288 |
| **GPCR** | | | |
| GAMMA-AMINOBUTYRIC ACID SIGNALING PATHWAY | 0.6150277 | 0.60434896 | 0 |
| POSITIVE REGULATION OF CYTOSOLIC CALCIUM ION CONCENTRATION | -0.5941538 | -0.57283056 | -0.57771325 |
| **Sensory system** | | | |
| DETECTION OF CHEMICAL STIMULUS INVOLVED IN SENSORY PERCEPTION OF SMELL | -0.5063316 | -0.52551603 | -0.47393885 |
| DETECTION OF CHEMICAL STIMULUS INVOLVED IN SENSORY PERCEPTION | -0.4913819 | -0.5083558 | -0.4614463 |
| DETECTION OF STIMULUS INVOLVED IN SENSORY PERCEPTION | -0.4738545 | -0.4874705 | -0.45169273 |
| ODORANT BINDING | -0.4926886 | -0.51260805 | -0.44146246 |
| OLFACTORY RECEPTOR ACTIVITY | -0.5063316 | -0.5255162 | -0.4739388 |
| SENSORY PERCEPTION OF CHEMICAL STIMULUS | -0.4697796 | -0.48142782 | -0.4410322 |
| SENSORY PERCEPTION OF SMELL | -0.4816634 | -0.4990263 | -0.45091802 |
| **Homeostasis** | | | |
| ESTABLISHMENT OF SKIN BARRIER | -0.6149926 | -0.5952358 | -0.6243971 |
| POSITIVE REGULATION OF CYTOSOLIC CALCIUM ION CONCENTRATION | -0.3052045 | -0.29191506 | 0 |
| REGULATION OF CYTOSOLIC CALCIUM ION CONCENTRATION | -0.2864533 | 0 | 0 |
| REGULATION OF WATER LOSS VIA SKIN | -0.5755355 | -0.5580616 | -0.5963385 |
