## Supplementary Table 3 for "Transcriptional profile of pyramidal neurons in chronic schizophrenia reveals lamina-specific dysfunction of neuronal immunity"

### Supplementary Table 3 Distinct pathways involved in superficial and deep neurons

| Pathway | SCZvsCtrl_D | SCZvsCtrl_S |
|---|---|---|
| POSTTRANSCRIPTIONAL REGULATION OF GENE EXPRESSION | 0.2971033 | 0 |
| REGULATION OF TRANSCRIPTION FROM RNA POLYMERASE II PROMOTER IN RESPONSE TO STRESS | 0.42602116 | 0 |
| REGULATION OF VIRAL TRANSCRIPTION | 0.45089537 | 0 |
| REPRESSING TRANSCRIPTION FACTOR BINDING | 0.4179329 | 0 |
| RNA POLYMERASE II REPRESSING TRANSCRIPTION FACTOR BINDING | 0 | 0.5335609 |
| TRANSCRIPTION-COUPLED NUCLEOTIDE-EXCISION REPAIR | 0.41460714 | 0 |
| VIRAL TRANSCRIPTION | 0 | -0.41784468 |
| CYTOPLASMIC TRANSLATION | 0.51095414 | 0 |
| MITOCHONDRIAL TRANSLATIONAL ELONGATION | 0 | 0.40039274 |
| TRANSLATION FACTOR ACTIVITY, RNA BINDING | 0.44332832 | 0 |
| TRANSLATION | 0.4067169 | 0 |
| TRANSLATIONAL INITIATION | 0.48899695 | 0 |
| PEPTIDYL-THREONINE PHOSPHORYLATION | 0 | 0.4143322 |
| POSITIVE REGULATION OF ERK1 AND ERK2 CASCADE | 0 | -0.33259064 |
| PROTEIN K11-LINKED UBIQUITINATION | 0.5870458 | 0 |
| PROTEIN OXIDATION | 0 | -0.67212313 |
| REGULATION OF ERK1 AND ERK2 CASCADE | -0.2856738 | 0 |
| REGULATION OF PROTEIN FOLDING | 0 | 0.74522674 |
| ATP METABOLIC PROCESS | 0.35959795 | 0 |
| ATPASE ACTIVITY | 0.2924602 | 0 |
| CELLULAR RESPIRATION | 0 | 0.33658525 |
| RESPIRATORY BURST | 0 | -0.666244 |
| INTRINSIC COMPONENT OF MITOCHONDRIAL MEMBRANE | 0.4374556 | 0 |
| MITOCHONDRIAL GENE EXPRESSION | 0 | 0.35573474 |
| MITOCHONDRIAL RIBOSOME | 0 | 0.3965855 |
| ESTABLISHMENT OF PROTEIN LOCALIZATION TO ENDOPLASMIC RETICULUM | 0.44145682 | 0 |
| PROTEIN LOCALIZATION TO ENDOPLASMIC RETICULUM | 0.42070618 | 0 |
| ESTABLISHMENT OF PROTEIN LOCALIZATION TO ORGANELLE | 0.29687834 | 0 |
| ORGANELLE INNER MEMBRANE | 0.29181665 | 0 |
| ACTIN CYTOSKELETON ORGANIZATION | 0.283129 | 0 |
| ESTABLISHMENT OR MAINTENANCE OF ACTIN CYTOSKELETON POLARITY | 0.7188997 | 0 |
| ESTABLISHMENT OR MAINTENANCE OF CYTOSKELETON POLARITY | 0.63730526 | 0 |
| REGULATION OF MITOTIC SPINDLE ASSEMBLY | 0.6320686 | 0 |
| NEURON PROJECTION CYTOPLASM | 0.40513325 | 0 |
| NEURON PROJECTION EXTENSION | 0.43597773 | 0 |
| NEURON SPINE | 0.3525331 | 0 |
| DENDRITIC SPINE | 0.3448264 | 0 |
| ASYMMETRIC SYNAPSE | 0.34774503 | 0 |
| GLUTAMATERGIC SYNAPSE | 0.29175848 | 0 |
| INTEGRAL COMPONENT OF POSTSYNAPTIC DENSITY MEMBRANE | 0.5126403 | 0 |
| INTRINSIC COMPONENT OF POSTSYNAPTIC SPECIALIZATION MEMBRANE | 0.43284112 | 0 |
| NEURON TO NEURON SYNAPSE | 0.34773207 | 0 |
| POSTSYNAPTIC DENSITY | 0.34773958 | 0 |
| POSTSYNAPTIC SPECIALIZATION | 0.33731592 | 0 |
| C-C CHEMOKINE RECEPTOR ACTIVITY | -0.6206737 | 0 |
| CCR CHEMOKINE RECEPTOR BINDING | 0 | -0.55911744 |
| GABA RECEPTOR COMPLEX | 0 | 0.61050135 |
| POSITIVE REGULATION OF T CELL RECEPTOR SIGNALING PATHWAY | -0.6455166 | 0 |
| SEROTONIN RECEPTOR SIGNALING PATHWAY | -0.48958847 | 0 |
| ACTIVE TRANSMEMBRANE TRANSPORTER ACTIVITY | -0.2867565 | 0 |
| AMMONIUM TRANSMEMBRANE TRANSPORTER ACTIVITY | -0.5454719 | 0 |
| ANION TRANSMEMBRANE TRANSPORT | -0.30251035 | 0 |
| ANION TRANSMEMBRANE TRANSPORTER ACTIVITY | -0.28741428 | 0 |
| CHOLESTEROL TRANSPORTER ACTIVITY | 0 | -0.6391872 |
| DICARBOXYLIC ACID TRANSMEMBRANE TRANSPORTER ACTIVITY | -0.50452155 | 0 |
| ICOSANOID TRANSPORT | -0.5237443 | 0 |
| LIPID TRANSPORTER ACTIVITY | 0 | -0.3867698 |
| MONOCARBOXYLIC ACID TRANSMEMBRANE TRANSPORTER ACTIVITY | -0.4570776 | 0 |

| | | |
|---|---|---|
| ORGANIC CATION TRANSMEMBRANE TRANSPORTER ACTIVITY | -0.55816746 | 0 |
| ORGANIC HYDROXY COMPOUND TRANSMEMBRANE TRANSPORTER ACTIVITY | -0.43071973 | 0 |
| STEROL TRANSPORTER ACTIVITY | 0 | -0.51992875 |
| NEUROTRANSMITTER:SODIUM SYMPORTER ACTIVITY | -0.5630056 | 0 |
| ARACHIDONIC ACID SECRETION | -0.5406368 | 0 |
| ICOSANOID SECRETION | -0.5228797 | 0 |
| REGULATION OF CYTOKINE SECRETION | -0.3155167 | 0 |
| REGULATION OF INTERLEUKIN-8 SECRETION | -0.5851303 | 0 |
| CYTOKINE ACTIVITY | -0.3044843 | 0 |
| CYTOKINE BINDING | -0.38870886 | 0 |
| CYTOKINE PRODUCTION | -0.34816307 | 0 |
| NEGATIVE REGULATION OF INTERLEUKIN-8 PRODUCTION | -0.62104726 | 0 |
| NEGATIVE REGULATION OF TUMOR NECROSIS FACTOR SUPERFAMILY CYTOKINE PRODUCTION | -0.43721634 | 0 |
| REGULATION OF INTERLEUKIN-8 PRODUCTION | -0.42007706 | 0 |
| LEUKOCYTE MIGRATION | -0.29767117 | 0 |
| LYMPHOCYTE CHEMOTAXIS | 0 | -0.5285425 |
| NEUTROPHIL CHEMOTAXIS | -0.39349398 | 0 |
| POSITIVE REGULATION OF LEUKOCYTE MIGRATION | 0 | -0.38535094 |
| POSITIVE REGULATION OF LYMPHOCYTE CHEMOTAXIS | 0 | -0.61327606 |
| POSITIVE REGULATION OF LYMPHOCYTE MIGRATION | 0 | -0.52468246 |
| POSITIVE REGULATION OF MONOCYTE CHEMOTAXIS | -0.61778307 | 0 |
| REGULATION OF LYMPHOCYTE CHEMOTAXIS | 0 | -0.5771987 |
| REGULATION OF NEUTROPHIL MIGRATION | 0 | -0.49605244 |
| REGULATION OF T CELL CHEMOTAXIS | 0 | -0.6250676 |
| GRANULOCYTE CHEMOTAXIS | -0.40334058 | 0 |
| GRANULOCYTE MIGRATION | 0 | -0.38834926 |
| C-C CHEMOKINE RECEPTOR ACTIVITY | -0.6206737 | 0 |
| CCR CHEMOKINE RECEPTOR BINDING | 0 | -0.55911744 |
| REGULATION OF LYMPHOCYTE PROLIFERATION | -0.3025959 | 0 |
| T CELL DIFFERENTIATION IN THYMUS | -0.43875417 | 0 |
| T CELL SELECTION | 0 | -0.50288785 |
| THYMIC T CELL SELECTION | -0.61653703 | 0 |
| POSITIVE REGULATION OF T CELL RECEPTOR SIGNALING PATHWAY | -0.6455166 | 0 |
| ANTIGEN PROCESSING AND PRESENTATION OF ENDOGENOUS ANTIGEN | 0 | -0.6458494 |
| ANTIGEN PROCESSING AND PRESENTATION OF EXOGENOUS ANTIGEN | 0.35665044 | 0 |
| ANTIGEN PROCESSING AND PRESENTATION OF EXOGENOUS PEPTIDE ANTIGEN VIA MHC CLASS II | 0.39800212 | 0 |
| ANTIGEN PROCESSING AND PRESENTATION OF EXOGENOUS PEPTIDE ANTIGEN | 0.3820784 | 0 |
| ANTIGEN PROCESSING AND PRESENTATION OF PEPTIDE OR POLYSACCHARIDE ANTIGEN VIA MHC CLASS II | 0.39800212 | 0 |
| IMMUNE RESPONSE-ACTIVATING SIGNAL TRANSDUCTION | 0 | -0.2844257 |
| IMMUNE RESPONSE-REGULATING SIGNALING PATHWAY | 0 | -0.2763649 |
| ANAPHASE-PROMOTING COMPLEX-DEPENDENT CATABOLIC PROCESS | 0.49522433 | 0 |
| MODIFICATION-DEPENDENT PROTEIN CATABOLIC PROCESS | 0.29391894 | 0 |
| PROTEASOMAL PROTEIN CATABOLIC PROCESS | 0.30749986 | 0 |
| PROTEASOMAL UBIQUITIN-INDEPENDENT PROTEIN CATABOLIC PROCESS | 0.6591135 | 0 |
| PROTEASOME-MEDIATED UBIQUITIN-DEPENDENT PROTEIN CATABOLIC PROCESS | 0.32165468 | 0 |
| PROTEIN DEUBIQUITINATION | 0.32102683 | 0 |
| PROTEIN MODIFICATION BY SMALL PROTEIN REMOVAL | 0.31958103 | 0 |
| UBIQUITIN-DEPENDENT PROTEIN CATABOLIC PROCESS | 0.2922979 | 0 |
| CELLULAR RESPONSE TO HYPOXIA | 0.3493245 | 0 |
| DEFENSE RESPONSE TO GRAM-POSITIVE BACTERIUM | -0.38255495 | 0 |
| REGULATION OF INFLAMMATORY RESPONSE | 0 | -0.3062746 |
| RESPONSE TO INTERFERON-GAMMA | 0 | -0.36389288 |
| GAMMA-AMINOBUTYRIC ACID SIGNALING PATHWAY | 0.60434896 | 0 |
| POSITIVE REGULATION OF CYTOSOLIC CALCIUM ION CONCENTRATION | -0.29191506 | 0 |

*0 represents insignificance
