## Supplementary Table 4 for "Transcriptional profile of pyramidal neurons in chronic schizophrenia reveals lamina-specific dysfunction of neuronal immunity"

**Supplementary Table 4 Antipsychotic treatment profiles**

| GEO code | Antipsychotics | Animal | Dose | Treatment length | Typical/Atypical |
|---|---|---|---|---|---|
| GSE48955 | Clozapine | Mouse | 3mg/kg | 1h | A |
| GSE48955 | Clozapine | Mouse | 3mg/kg | 2h | A |
| GSE48955 | Clozapine | Mouse | 3mg/kg | 4h | A |
| GSE48955 | Clozapine | Mouse | 3mg/kg | 8h | A |
| GSE48955 | Risperidone | Mouse | 0.5mg/kg | 1h | T |
| GSE48955 | Risperidone | Mouse | 0.5mg/kg | 2h | T |
| GSE48955 | Risperidone | Mouse | 0.5mg/kg | 4h | T |
| GSE48955 | Risperidone | Mouse | 0.5mg/kg | 8h | T |
| GSE48955 | Haloperidol | Mouse | 1mg/kg | 1h | T |
| GSE48955 | Haloperidol | Mouse | 1mg/kg | 2h | T |
| GSE48955 | Haloperidol | Mouse | 1mg/kg | 4h | T |
| GSE48955 | Haloperidol | Mouse | 1mg/kg | 8h | T |
| GSE48955 | Clozapine | Mouse | 3mg/kg | 1h | A |
| GSE48955 | Clozapine | Mouse | 3mg/kg | 2h | A |
| GSE48955 | Clozapine | Mouse | 3mg/kg | 4h | A |
| GSE48955 | Clozapine | Mouse | 3mg/kg | 8h | A |
| GSE48955 | Risperidone | Mouse | 0.5mg/kg | 1h | T |
| GSE48955 | Risperidone | Mouse | 0.5mg/kg | 2h | T |
| GSE48955 | Risperidone | Mouse | 0.5mg/kg | 4h | T |
| GSE48955 | Risperidone | Mouse | 0.5mg/kg | 8h | T |
| GSE33822 | Clozapine | Mouse | 4mg/kg | 3h | A |
| GSE33822 | Clozapine | Mouse | 4mg/kg | 3h | A |
| GSE33822 | Clozapine | Mouse | 4mg/kg | 3h | A |
| GSE4031 | Clozapine | Rat | Low | Unknown | A |
| GSE4031 | Clozapine | Rat | Medium | Unknown | A |
| GSE4031 | Clozapine | Rat | High | Unknown | A |
| GSE4031 | Haloperidol | Rat | Low | Unknown | T |
| GSE4031 | Haloperidol | Rat | Medium | Unknown | T |
| GSE4031 | Haloperidol | Rat | High | Unknown | T |
| GSE66275 | Haloperidol | Rat | 0.25mg/kg | 21d | T |
| GSE66275 | Haloperidol | Rat | 0.25mg/kg | 21d | T |
| GSE66275 | Haloperidol | Rat | 0.25mg/kg | 21d | T |
| GSE66275 | Risperidone | Rat | 5mg/kg | 21d | A |
| GSE66275 | Risperidone | Rat | 5mg/kg | 21d | A |
| GSE66275 | Risperidone | Rat | 5mg/kg | 21d | A |
| GSE45229 | Haloperidol | Mouse | 0.3 mg/kg | 28d | T |
| GSE45229 | Haloperidol | Mouse | 1 mg/kg | 28d | T |
| GSE45229 | Quetiapine | Mouse | 10 mg/kg | 28d | A |
| GSE45229 | Quetiapine | Mouse | 100 mg/kg | 28d | A |
| GSE45229 | Haloperidol | Mouse | 0.3 mg/kg | 28d | T |
| GSE45229 | Haloperidol | Mouse | 1 mg/kg | 28d | T |
| GSE45229 | Quetiapine | Mouse | 10 mg/kg | 28d | A |
| GSE45229 | Quetiapine | Mouse | 100 mg/kg | 28d | A |
| GSE125325 | Quetiapine | Rat oligodendrocyte precursor cells | 1 μM | 48h | A |
| GSE125325 | Quetiapine | Rat oligodendrocyte precursor cells | 1 μM | 96h | A |
| GSE6467 | Clozapine | Mouse | 12mg/kg | 12wk | A |
| GSE6467 | Haloperidol | Mouse | 1.6mg/kg | 12wk | T |
| GSE6511 | Clozapine | Mouse | 12mg/kg | 4wk | A |
| GSE6511 | Haloperidol | Mouse | 1.6mg/kg | 4wk | T |
| GSE89873 | Haloperidol | Rat | 25μM | 2h | T |
| GSE677 | Haloperidol | Mouse | 3mg/kg | 30d | T |
